# Relating enhancer genetic variation across mammals to complex phenotypes using machine learning

**DOI:** 10.1101/2022.08.26.505436

**Authors:** Irene M. Kaplow, Alyssa J. Lawler, Daniel E. Schäffer, Chaitanya Srinivasan, Morgan E. Wirthlin, BaDoi N. Phan, Xiaomeng Zhang, Kathleen Foley, Kavya Prasad, Ashley R. Brown, Zoonomia Consortium, Wynn K. Meyer, Andreas R. Pfenning

## Abstract

Protein-coding differences between mammals often fail to explain phenotypic diversity, suggesting involvement of enhancers, often rapidly evolving regions that regulate gene expression. Identifying associations between enhancers and phenotypes is challenging because enhancer activity is context-dependent and may be conserved without much sequence conservation. We developed TACIT (Tissue-Aware Conservation Inference Toolkit) to associate open chromatin regions (OCRs) with phenotypes using predictions in hundreds of mammalian genomes from machine learning models trained to learn tissue-specific regulatory codes. Applying TACIT for motor cortex and parvalbumin-positive interneurons to neurological phenotypes revealed dozens of new OCR-phenotype associations. Many associated OCRs were near relevant genes, including brain size-associated OCRs near genes mutated in microcephaly or macrocephaly. Our work creates a forward genomics foundation for identifying candidate enhancers associated with phenotype evolution.

**One Sentence Summary:** A new machine learning-based approach associates enhancers with the evolution of brain size and behavior across mammals.

## INTRODUCTION

Much of the phenotypic diversity that exists across vertebrates is thought to have arisen from differences in how genes are expressed (*1*). Variation in phenotypes like vocal learning (*2*) and longevity (*3*) has been linked to patterns of gene expression within some of the most relevant brain regions and tissues, respectively. Thus, many genetic differences associated with the evolution of these, and other, complex phenotypes are likely in enhancers, distal *cis*-regulatory genomic elements that are bound by transcription factor (TF) proteins that regulate the expression of associated genes, often through cell type-specific activation (*4, 5*). For example, limblessness in snakes is associated with sequence divergence and activity loss in a critical enhancer near the *Sonic hedgehog* gene (*6*), and mutations in orthologs of this enhancer are associated with polydactyly in humans, mice, and cats (*7, 8*). Enhancer evolution has been found to be associated with a number of other complex phenotypes, including eyesight loss (*9*) as well as whisker, penile spine, and brain growth (*10*).

Recent advances facilitate identifying relationships between enhancer activity and phenotype evolution. Community genome sequencing efforts such as the Zoonomia Project have constructed assemblies for hundreds of species from diverse mammalian clades (*11*). Cactus multi-species whole-genome alignments and tools for extracting orthologs have vastly improved ortholog mapping for non-coding genomic regions (*12–14*). In addition, new phylogeny-aware statistical methods have been developed for identifying factors associated with the evolution of phenotypes (*15, 16*).

Despite these successes, identifying enhancer-phenotype relationships is still a major challenge. Widely used methods to identify conservation and convergent evolution across orthologous genome sequences measure the extent to which the nucleotides within a given region align across species (*17–19*). While these approaches have led to some exciting findings (*9, 20*), many enhancer sequences and transcription factor binding sites are under less sequence constraint than promoter and gene sequences (*21, 22*). In fact, recent studies have shown that sequence conservation is not required for activity conservation at enhancer orthologs (*23, 24*) and can occur when enhancer activity is not conserved in a tissue of interest (*25*), so nucleotide sequence conservation at enhancers is sometimes an insufficient proxy for enhancer activity conservation.

Here we present a new method for identifying enhancer-phenotype associations, in which we trace enhancer activity evolution using predicted open chromatin in a tissue or cell type of interest as a proxy for enhancer function. Previously, we and others have demonstrated that the sequence patterns associated with enhancer activity in multiple tissues are highly conserved across mammals by showing that machine learning models that use DNA sequence to predict enhancer activity in a tissue of interest in one species can accurately predict clade-specific and tissue-specific enhancer activity in species from different mammalian clades (*25,27–29*). We integrate machine learning-based predictions of enhancer function with other comparative genomics advances (*11, 15, 16*) in a new framework called the Tissue-Aware Conservation Inference Toolkit (TACIT) for identifying candidate enhancers associated with the evolution of phenotypes. We use sequences underlying open chromatin regions (OCRs) from a small number of species in a tissue or cell type of interest to train convolutional neural networks (CNNs) that predict the probability of OCR ortholog open chromatin in those tissues/cell types at the orthologous sequences in up to 222 mammalian genomes (*11*). We then use these predictions to link OCRs to specific mammalian phenotypes while accounting for phylogeny (**Fig. 1**). We applied our approach to multiple phenotypes, including brain size, solitary and group living, and vocal learning, and identified both motor cortex tissue and motor cortex parvalbumin-positive (PV+) interneuron OCRs associated with these phenotypes that are near relevant genes. Our approach can be applied to any phenotype with open chromatin data available from a relevant tissue or cell type in at least two species. It is therefore broadly applicable to a variety of tissue, phenotype, and species combinations.

**Figure 1:**
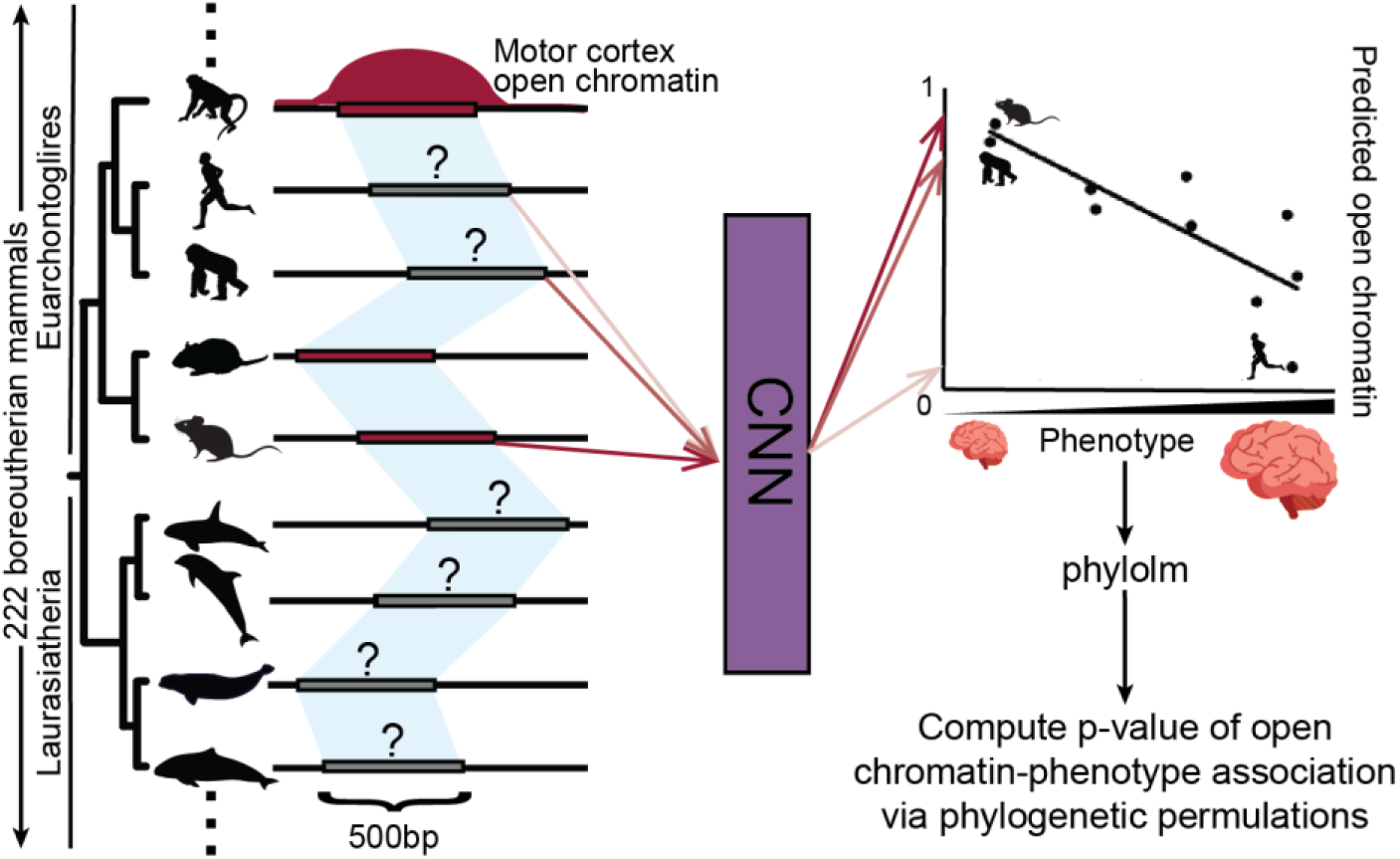
Overview of TACIT. We train a CNN using sequences underlying OCRs and non-OCRs to predict open chromatin in a tissue or cell type of interest and then use the CNN to predict open chromatin in that tissue or cell type in hundreds of genomes from Zoonomia. We associate our predictions with phenotypes using phylolm and then quantify the significance of the association using an empirical p-value from phylogenetic permutations. Animal silhouettes were made by Michael Keesey, Daniel Jaron, Ryan Cupo, Steven Traver, and Chris Huh (license: https://creativecommons.org/licenses/by-sa/3.0/); were downloaded from PhyloPic; and were not modified.

## RESULTS

### Convolutional neural networks accurately predict open chromatin status of OCR orthologs

We applied TACIT to two tissues with open chromatin data from more than two species – motor cortex and liver – as well as a tissue and a cell type with data from only two species – retina and motor cortex PV+ interneurons. We used OCRs instead of other enhancer activity measures, such as H3K27ac ChIP-seq regions, because OCRs tend to have a concentration of TF motifs near their summits and be hundreds instead of thousands of base pairs long, allowing our model to focus on sequences likely to be involved in enhancer activity and allowing us to easily map regions in species whose assemblies have short scaffolds (*14*). We chose tissues and cell types that would demonstrate specificity in dissimilar tissues (brain versus liver) and have relationships with complex phenotypes of interest, including brain size, social behavior, and vocal learning. For tissues with more than two species, we trained CNNs to predict whether a region is an OCR or a non-OCR ortholog of an OCR, as described in our previous work (*25*).

Since we are the first to train machine learning models for open chromatin prediction in motor cortex (we and others have shown that the liver regulatory code is conserved across species (*25, 27*)), we first trained CNNs using only house mouse sequences and found that the CNNs successfully predicted clade-specific OCRs and non-OCRs (high “lineage-specific OCR accuracy,” AUC > 0.70 and AUPRC/NPV-Spec. > 0.65 for all metrics) as well as tissue-specific OCRs and non-OCRs (high “tissue-specific OCR accuracy,” AUC > 0.65 and AUPRC/NPV-Spec. > fraction of examples in smaller class for all metrics); in addition, when comparing average OCR ortholog predictions across species, predictions had the expected negative correlation with distance from the species in which the OCRs were assayed (high “phylogeny-matching correlations,” mean Pearson correlation < -0.70 and mean Spearman correlation < -0.45) (**Figs. S1A,D,G,J,M,P, Table S1**) (*25*). We next trained multi-species CNNs for motor cortex and liver using all of our data – *Mus musculus* (Glires clade), *Macaca mulatta* (Euarchonta clade), and *Rattus norvegicus* (Glires clade) for both tissues as well as *Rousettus aegyptiacus* (Laurasiatheria clade) for motor cortex and *Bos taurus* (Laurasiatheria clade) and *Sus scrofa* (Laurasiatheria clade) for liver – and found that the models achieved high lineage- and tissue-specific OCR accuracy (AUC > 0.8, AUPRC/NPV-Spec. > fraction of examples in smaller class for all metrics) as well as phylogeny-matching correlations (mean Pearson correlation < -0.95 and mean Spearman correlation < -0.75) (**Fig. 2**, **Figs. S2A,D,G, Fig. S3, Tables S2-3**). We then used the multi-species motor cortex CNN to make predictions at motor cortex OCR orthologs in 222 diverse boreoeutherian mammal genomes from Zoonomia, where we limited ourselves to boreoeutherians because we did not have open chromatin data from species in other clades. To further evaluate the reliability of our predictions, we clustered the species hierarchically with predictions as features and found that the cluster hierarchy was similar to the phylogenetic tree, with all but a few species clustering correctly by clade (**Fig. S4, Supplementary Text**) (*26*).

**Figure 2:**
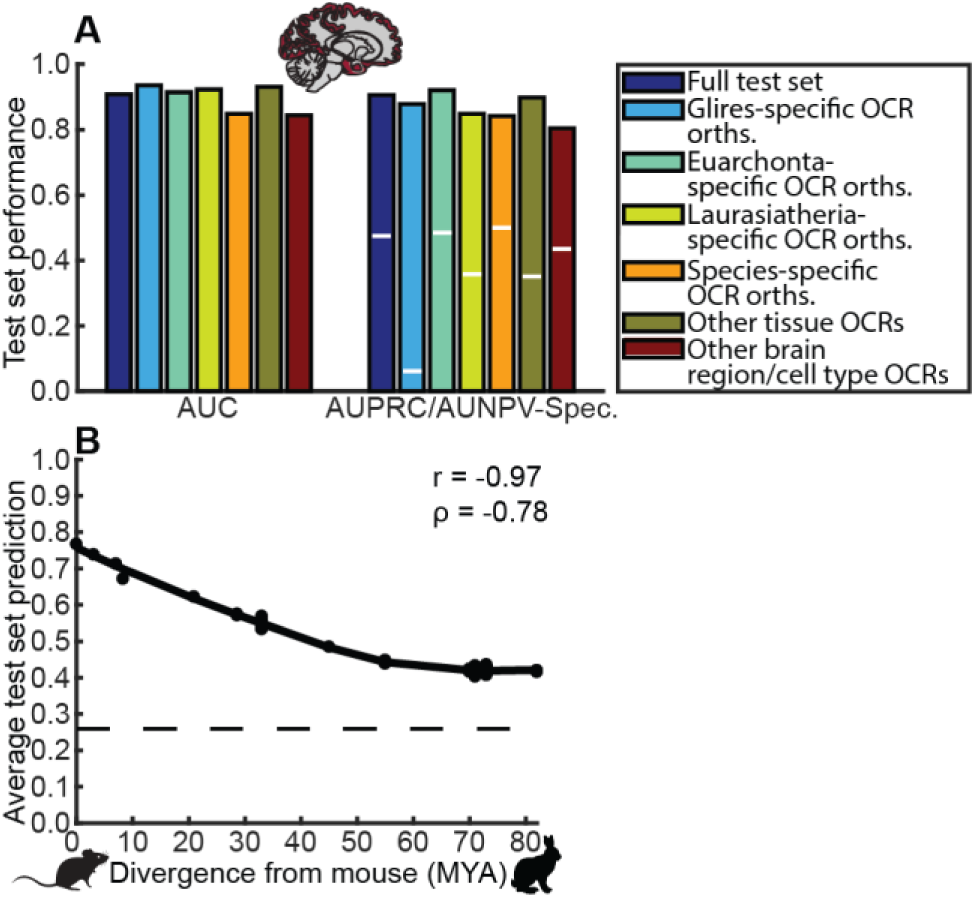
Motor cortex multi-species CNN performance. **A** shows the area under the ROC curve (AUROC) and the area under the precision-recall (if more negatives than positives)/negative predictive value-specificity (if more positives than negatives) curve (AUPRC/NPV-Spec.) for the full test set, clade-specific OCRs and non-OCRs, and shared versus tissue/brain region-specific OCRs and non-OCRs for the multi-species motor cortex CNNs. **B** shows the negative relationship between the average house mouse OCR ortholog multi-species motor cortex open chromatin predicted probabilities for Glires species and the millions of years ago (MYA) when each species diverged from house mouse.

Since no previous study has trained PV+ interneuron or retinal enhancer activity prediction models for predicting enhancer activity in species not used for training (*25,27–29*), we needed to investigate whether the PV+ interneuron and retinal regulatory codes are sufficiently conserved for accurately predicting open chromatin of OCR orthologs. We did this by running motif discovery on open chromatin datasets from each species for which data was available. For each of PV+ interneurons and retina, we found motifs for many of the same TFs in both species, and some of these TFs are known to be involved in PV+ interneurons and retina, respectively (**Supplementary Text, Supplementary Website**) (*26*).

Because we had PV+ interneuron and retina data from only two species – *Mus musculus* and *Homo sapiens* (Euarchonta clade) – we did not have sufficient non-OCR orthologs of OCRs to train CNNs, so we developed a new approach to constructing negative sets for these cases: We combined a large number of random regions of the genome with the same G/C-content as the positives with OCRs from other cell types or tissues, two negative sets that provided adequate performance for all of our metrics in our previous work (**Methods**) (*25*). To ensure that CNNs could make accurate predictions in species not used for training in our tissues and cell types, we first trained CNNs using only house mouse sequences and found that they achieved high lineage-specific OCR accuracy (AUC > 0.85 and AUPRC/NPV-Spec. > 0.60) as well as phylogeny-matching correlations (mean Pearson correlation < - 0.65, mean Spearman correlation < -0.40 for retina and PV+ interneurons) for house mouse sequences (**Figs. S1B,C,E,F,H,I,K,L,N,O,Q,R, Tables S4-5**). The PV+ interneuron CNNs also achieved strong performance on human sequences (AUC > 0.70 and AUPRC/NPV-Spec. > fraction of examples in minority class for all criteria), where no human sequences were used in training as well as high tissue-specific OCR accuracy (AUC > 0.75 and AUPRC/NPV-Spec. > fraction of examples in minority class for all criteria), while the house mouse-trained retina CNNs did not work as well on human-specific OCRs and non-OCRs and liver non-retina OCRs. We then trained CNNs using sequences from both house mouse and human, and both the PV+ and retina CNNs achieved strong performance for all criteria (AUC > 0.70 and AUPRC/NPV-Spec. > fraction of examples in minority class for all criteria, mean Pearson correlation < -0.60, mean Spearman correlation < -0.40) (**Figs. 3A-D**, **Figs. S2B,C,E,F,H,I, Tables S6- 7**).

**Figure 3:**
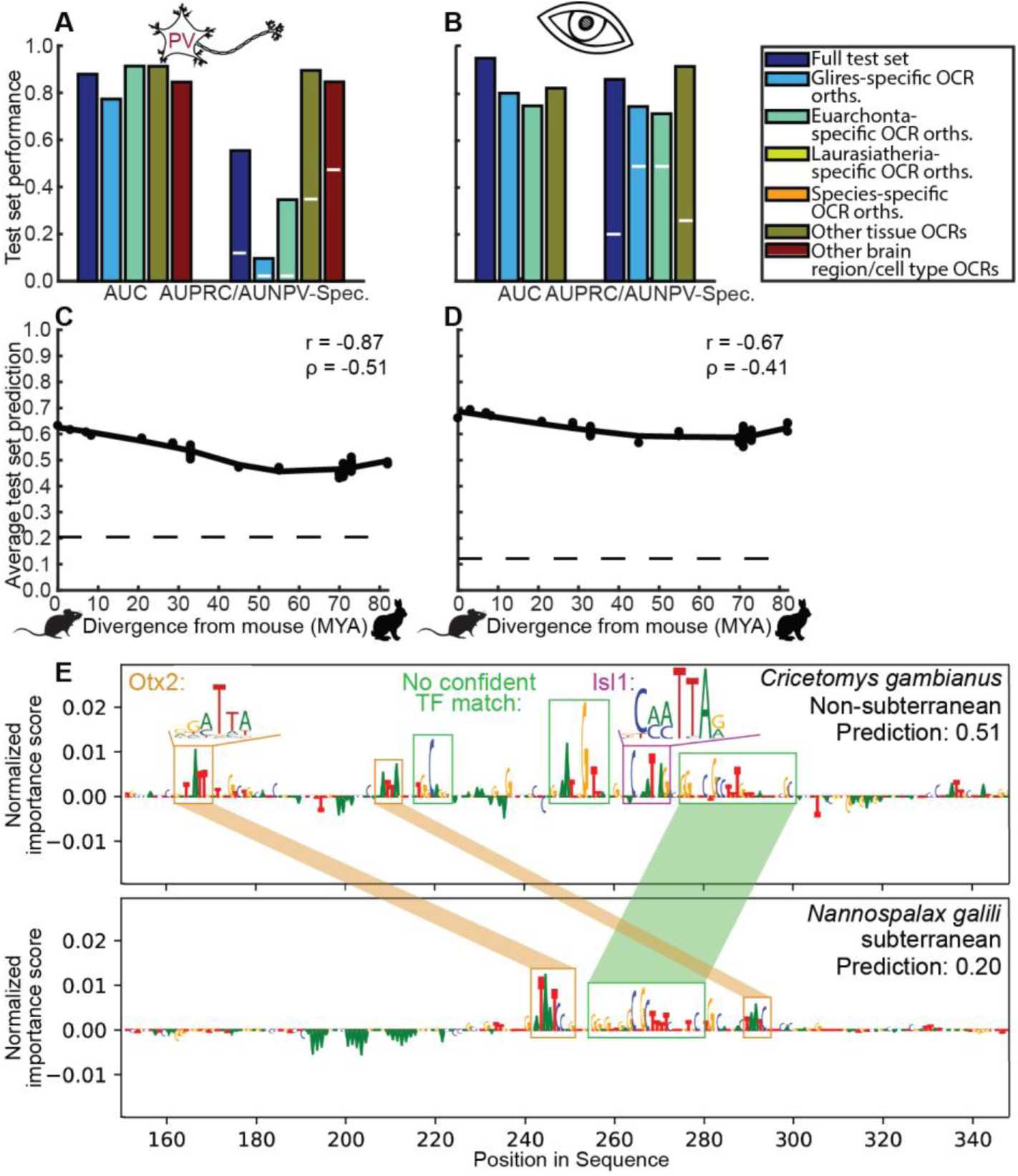
PV+ interneuron and retina multi-species CNN performance. (**A-B**) show the AUROC and the AUPRC/NPV-Spec. for the full test set, clade- specific OCRs and non-OCRs, and shared versus tissue/cell type-specific OCRs and non-OCRs for multi-species PV+ interneuron (**A**) and retina (**B**) CNNs. (**C-D**) show the negative relationship between the average house mouse OCR ortholog multi- species PV+ interneuron (**C**) and retina (**D**) CNN predictions for Glires species and the MYA when each species diverged from house mouse. E shows the multi-species retina model normalized importance scores for each position in the summit +/- 100bp of an OCR near *OTX2* that was previously shown to have a higher relative evolutionary rate in subterranean mammals. Orange boxes mark matches to the house mouse Otx2 motif, the magenta box marks the match to the house mouse Isl1 motif, and green boxes mark regions with high importance scores that do not match any known TF motif. Motifs were downloaded from CIS-BP (*86*) and visualized using meme2images from the MEME suite (*87*). No nucleotides in either ortholog outside these central 200 base pairs had a normalized importance score with absolute value greater than one.

To evaluate if our bulk tissue models were learning sequences relevant to the tissues in which they were trained, we interpreted what they had learned (**Methods**). Specifically, we computed the CNNs’ per-nucleotide importance scores, which indicate the extent to which the CNN prioritizes the presence or absence of each nucleotide at each position (*30, 31*). We found that our CNNs seemed to have learned sequence patterns that are similar to motifs of TFs that are known to be involved in motor cortex and liver, such as MEF2C for motor cortex (*32, 33*) and HNF4A (*34, 35*) for liver, as well as sequence patterns that do not match any known TF motif (**Supplementary Text**, **Figs. S5-7**) (*26*). We then examined a specific retina OCR near the retina TF *Otx2*, where the OCR’s orthologs in subterranean mammals were previously shown to have a faster relative evolutionary rate than its orthologs in other mammals (*9*). This OCR’s ortholog in *Nannospalax galili*, a subterranean mole-rat, was confidently predicted to be closed, while its ortholog in a non-subterranean pouched rat, *Cricetomys gambianus*, the most closely related mammal in Zoonomia that never lives underground (diverged ∼45 MYA (*36*)), was predicted to be open. Both of these OCR orthologs contained two motifs for Otx2 as well as a third motif that could not be easily interpreted with high importance scores. In addition to those important sequences, the *Cricetomys gambianus* ortholog had a high importance score for the motif for Isl1, a transcription factor involved in the development of bipolar and cholinergic amacrine cells of the retina (*37*). There were also two additional sequences with high importance scores unique to *Cricetomys gambianus* relative to *Nannospalax galili* that did not match any known TF motif, demonstrating the value of using a modeling strategy that does not require featurizing the sequence based on known information (**Fig. 3E**).

From the four cross-species OCR datasets of interest (motor cortex, liver, PV+ interneuron, and retina), we identify 50,942,699 total orthologous regions across 222 Boreoeutherian mammals from 402,880 total OCRs. Relative to human OCR annotations and phyloP annotations alone, we find that these predictions can provide a substantial boost for interpreting human disease-associated loci, with greater tissue- and cell type specificity. For example, in our other work, we found that human orthologs of regions predicted to have conserved motor cortex open chromatin are enriched for overlapping SNPs associated with schizophrenia, while human orthologs of regions predicted to have conserved liver open chromatin are enriched for overlapping SNPs associated with cholesterol-related traits (*38, 39*). These results demonstrate the power of TACIT to identify functionally relevant patterns of conservation.

### Applying TACIT to mammalian phenotypes

#### A framework for associating predicted open chromatin with phenotypes

Having trained models to predict open chromatin status of OCR orthologs in four tissues and cell types – motor cortex, liver, retina, and PV+ interneurons within the motor cortex – we identified individual OCRs whose predicted open chromatin across species is associated with phenotypes (**Fig. 1**). We applied the phylolm and phyloglm methods (*15*) for continuous and binary traits, respectively. These methods are modifications of phylogenetic generalized least squares (*40, 41*) designed for faster performance. We used them to test for a relationship between one OCR ortholog’s open chromatin predictions across species and phenotype annotations across species that cannot be explained by the species phylogeny alone. To minimize false positives, we implemented phylogenetic permulations (*16*), enabling us to evaluate the significance of each OCR-phenotype relationship against a background distribution of shuffled phenotypes with similar phylogenetic structures (**Materials and Methods**).

#### TACIT identifies motor cortex and PV+ interneuron OCRs associated with the evolution of brain size

We used TACIT to identify motor cortex OCRs whose predicted open chromatin across mammals is significantly associated with brain size, a complex trait with great diversity across mammals that is thought to underlie human cognitive ability (*42*). As brain size scales with body size, we used the brain size residual (brain mass minus the predicted value of brain mass from a regression on body mass), which we obtained for 158 boreoeutherian mammals (*43, 44*). Before applying TACIT, we investigated whether there are proteins whose relative evolutionary rates (*19*) are associated with the evolution of brain size residual. We did not find any proteins with a significant association after RERconverge’s default multiple hypothesis correction (corrected p ≥ 0.05 for all genes) (*19, 45*), which corroborates evidence that the top decile of TFs with the highest fraction of conserved base pairs tend to be enriched for embryonic development and brain function (PhyloP ≥ 2.241, FDR < 5%) (*39*) and previous work suggesting that enhancer loss drove the evolution of human-specific patterns in brain growth (*10*). In contrast, using TACIT, we found 34 motor cortex OCRs with a significant association with brain size residual after false discovery rate correction (α=0.05). We then examined all genes near (TSSs within 1Mb) those OCRs. Of the associated OCRs, 29 are near genes whose corresponding proteins play important roles in brain development, and 6 are near genes whose corresponding proteins are involved in brain tumor growth (**Table S8**). While many of these genes may influence brain size during development, the OCRs that regulate them might continue to be open during adulthood. This would be consistent with recent evidence that neural progenitors are responsible for the evolution of brain size in the great apes (*46*).

Of the 29 brain size residual-associated OCRs near brain development genes, 23 are near genes with mutations that cause neurological disorders, including 8 OCRs near genes in which mutations have been reported to cause microcephaly or macrocephaly (**Table S8, Figs. S8A-H**) (*47*). Furthermore, we found that the p-values of all motor cortex OCRs whose human orthologs are near (in hg38 coordinates) genes mutated in microcephaly or macrocephaly have a significantly lower distribution than the p-values of other motor cortex OCRs with human orthologs (p=0.0073, 1-sided Wilcoxon rank-sum test).

We identified two OCRs near *SATB1* — a gene with both microcephaly- and macrocephaly-associated mutations (*48*) — whose motor cortex predicted open chromatin status is significantly associated with brain size residual (**Fig. 4A-B, Figs. S8D**,**H**). For both of these associations, predicted open chromatin is associated with small brain size residual. The OCRs’ coordinates in the genomes in which they were initially identified are chr17:52351209-52351928 (mm10) and chr2:174466184- 174466517 (rheMac8). They are each about 500kb from the TSS of the gene, where one is upstream and the other is downstream. Neither OCR is near any other gene with a known connection to brain development; *Satb1*/*SATB1* is the second-closest gene to each, and the closer genes, *Kcnh8* and *TBC1D5*, each have known roles outside of brain growth (*49, 50*). The associations seem to be driven in large part by, respectively, cetaceans (**Fig. 4A**) and great apes (**Fig. 4B**), both of which have a large variation in brain size (*51*). In particular, the latter OCR is predicted to be active in all great apes except for humans, the great ape with the largest brain size residual. Interestingly, the reported case of *SATB1*-associated macrocephaly at birth was caused by a mutation that disrupts a large portion of the protein product, while microcephaly was usually reported with *SATB1* missense mutations (*48*). This pattern is consistent with the significant negative associations between predicted open chromatin and brain size residual, assuming that the OCRs we identified positively regulate the expression of *SATB1*.

**Figure 4:**
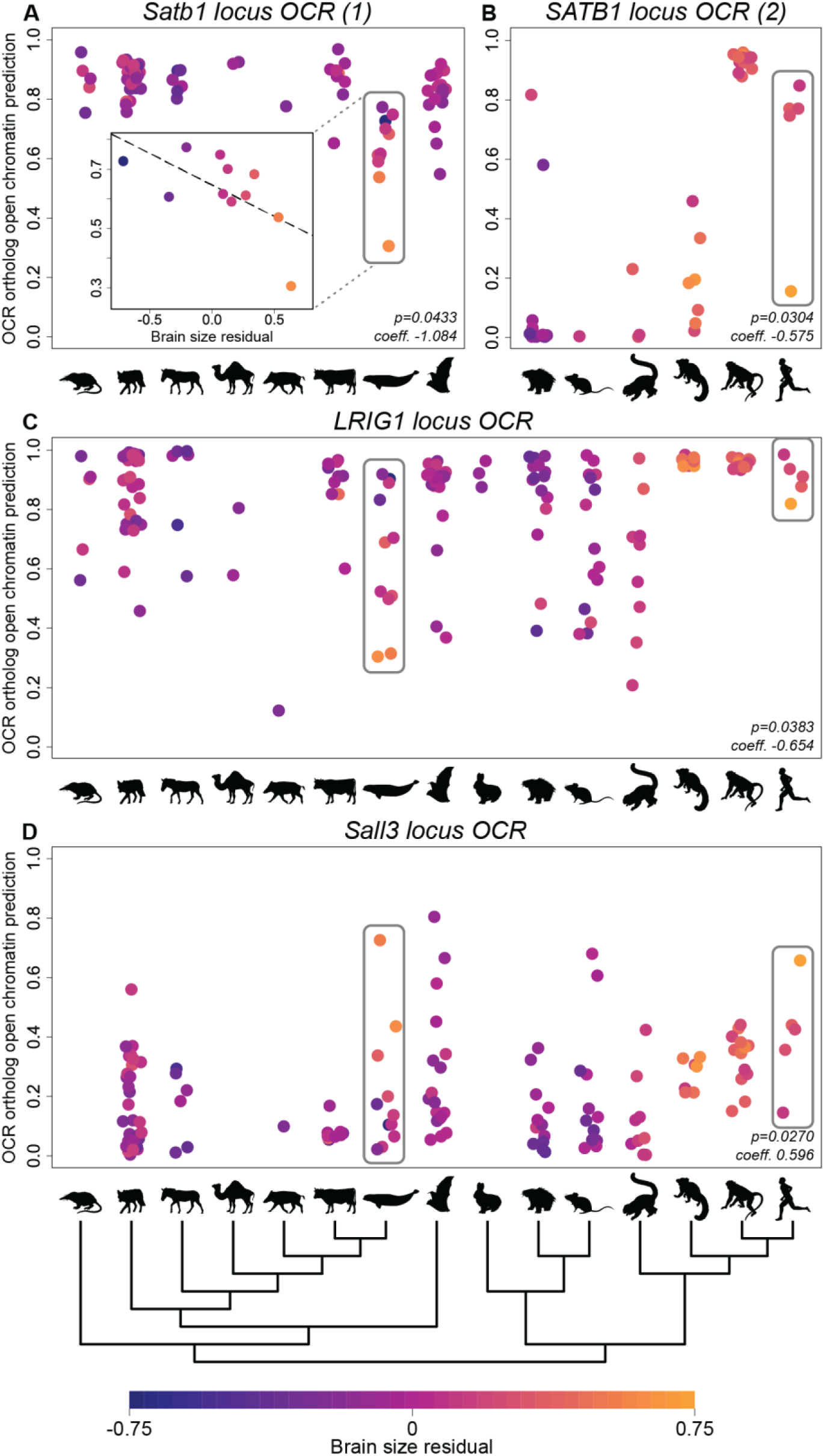
Examples of associations between predicted motor cortex OCR ortholog open chromatin and brain size residual. (**A-B**) highlight the negative association between predicted motor cortex open chromatin and brain size residual of orthologs of two motor cortex OCRs in the *SATB1* locus, chr17:52351209-52351928 (mm10) and chr2:174466184- 174466517 (rheMac8), within Laurasiatheria and Euarchontoglires, respectively. The latter OCR has no orthologs in Lagomorpha, which is omitted from panel (**B**). Boreoeutherian mammal-wide panels are shown in Fig. S9. (**C**) highlights the negative association of orthologs of a motor cortex OCR in the *LRIG1* locus, chr15:40082805- 40083380 (mm10). (**D**) highlights the positive association of orthologs of a motor cortex OCR in the *Sall3* locus, chr18:81802310- 81802951 (mm10). Each point represents one ortholog; they are grouped along the x-axis of each panel by clade as shown by the tree below. The clades and example species are listed in Table S10. The hominoid and cetacean clades are highlighted by gray boxes in each panel.

We identified another OCR, chr2:75345159-75346046 (rheMac8), whose predicted motor cortex open chromatin also has a strong negative association with brain size residual in cetaceans (**Fig. 4C**). The closest gene to this OCR is *LRIG1*, which is about 250kb from the OCR. *LRIG1* slows and delays the differentiation of neural stem cells (*52, 53*). While this OCR is also near other genes, none of those genes have a known role in brain size.

Points are colored by brain size residual following the scale at the bottom. The permulations p- value after Benjamini-Hotchberg correction and the coefficient on the predicted open chromatin returned by phylolm are shown in the lower right of each panel.

Also among the OCRs we identified near brain development genes is an OCR, chr18:81802310- 81802951 (mm10), about 800kb from the gene *Sall3*. Sall3 is the fourth-closest gene to this OCR, and one closer gene, *Mbp*, does have a connection to brain development (*54*). Hi-C from adult human cortex (*55*) shows that the bin containing the human ortholog of this OCR is close to *SALL3* in 3D space (p=2.3 X 10^-11^, **Table S8**) but is not close to MBP (*p*=1). This OCR displays a positive association with brain size residual both overall and within mammalian clades with especially large variations in brain size, including the great apes and cetaceans (**Fig. 4D**). *Sall3* is a member of the spalt-like family of transcription factors, which are important in development (*56*). Although a specific role of *Sall3* in motor cortex has not been described, there is evidence that *Sall3* regulates the maturation of neurons in other regions of the brain (*57, 58*), and *Sall3* is expressed in developing motor neurons (*58*) and human cerebral cortex (*59*).

We extended our framework to establish Cell-TACIT, a version of TACIT that identifies OCRs in specific cell types (*60, 61*) whose open chromatin predictions are associated with a phenotype of interest. We used Cell-TACIT for PV+ interneurons within the motor cortex to identify such OCRs whose predicted activity across Euarchontoglires is significantly associated with brain size residual. PV+ interneurons are a minority population, representing roughly 4 - 8% of neurons and 2 - 4 % of the total cell population in the mouse cortex (*62*) yet are critical in cortical microcircuits and human brain disorders like schizophrenia (*63, 64*). Given this sparsity, our bulk motor cortex open chromatin data may not capture OCRs that are specific to PV+ interneurons. In fact, about 30% of mouse PV+ OCRs do not overlap any bulk motor cortex OCRs, including non-reproducible peaks. We identified 13 OCRs whose predicted open chromatin in PV+ interneurons is associated with species’ brain size residuals after false discovery rate correction (α=0.05) (**Table S9**), 11 of which are house mouse OCRs for which predicted open chromatin is associated with having a smaller brain size residual.

We identified three PV+ interneuron OCRs that are significantly negatively associated with brain size residual and are within 1Mb of a gene that is mutated in macrocephaly or microcephaly (**Table S9**, **Figs. S8I-K**). Two of those OCRs — chr13:114757413-114757913 (mm10) and chr13:114793237-114793737 (mm10) — are respectively about 60kb and 25kb from the *Mocs2* gene. Both have strong associations with brain size residual within Euarchonta (primates and their closest relatives), especially Hominoidea, and the first also has some association within Glires (rodents and their closest relatives) (**Fig. 5A-B**, respectively). *Mocs2* is one of four genes involved in Molybdenum cofactor biosynthesis (*65*). Molybdenum cofactor deficiency (MoCD) in humans is a rare, fatal disease marked by intractable seizures, hypoxia, and microcephaly (*66*). We also identified an OCR, chr1:95762160-95762660 (mm10), that is about 100kb away from the gene *St8sia4*, which is important for the development and density of interneurons — including PV+ interneurons — in the cortex (*67, 68*).

**Figure 5:**
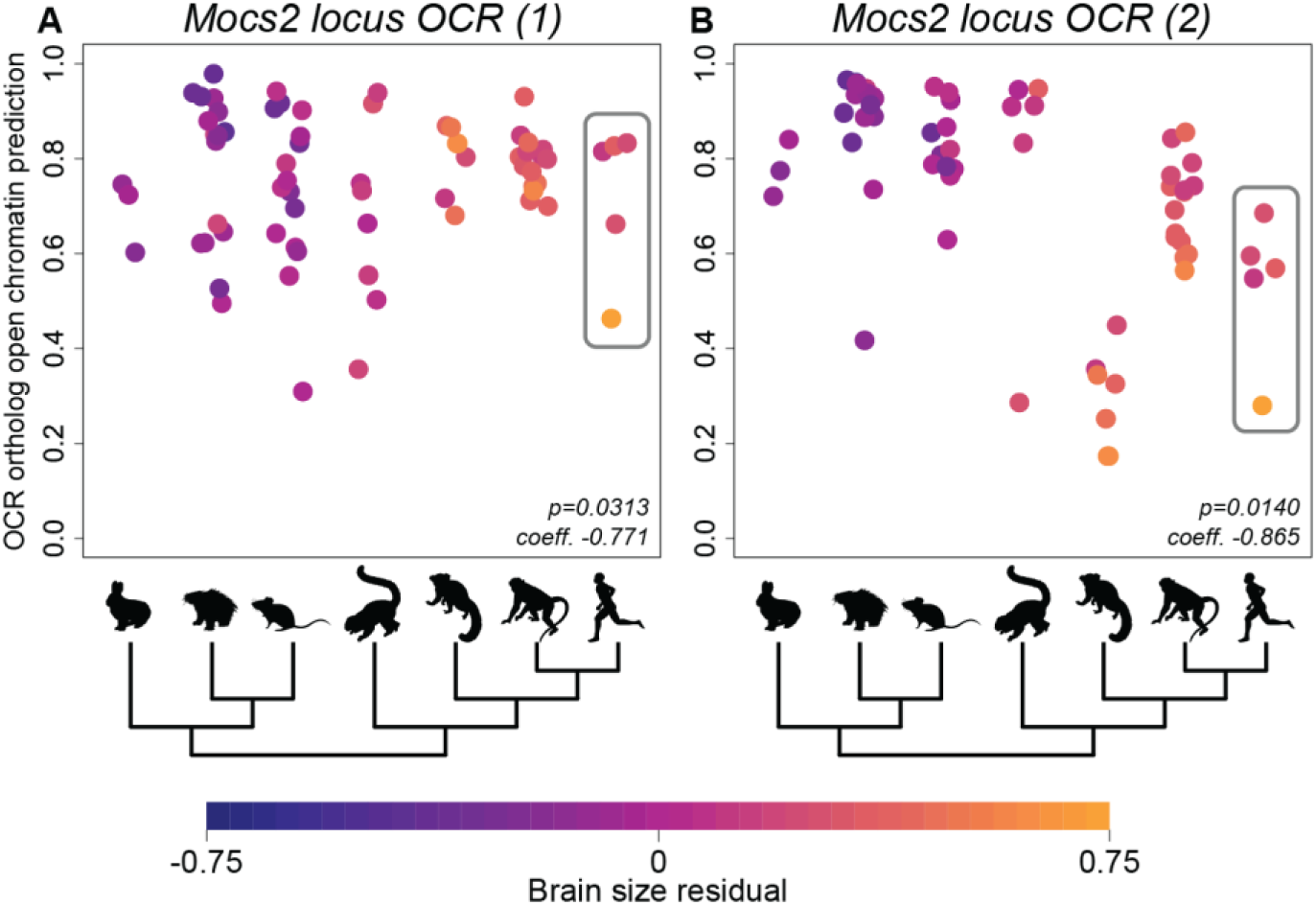
Examples of associations between predicted PV+ interneuron OCR ortholog open chromatin and brain size residual. (**A-B**) highlight the negative association within Euarchontoglires between predicted PV+ interneuron open chromatin and brain size residual of orthologs of two PV+ interneuron OCRs in the *Mocs2* locus, chr13:114757413-114757913 (mm10) and chr13:114793237-114793737 (mm10). Each point represents one ortholog; they are grouped along the x-axis of each panel by clade as shown by the tree below. The clades and example species are listed in Table S10. The hominoid clade is highlighted by a gray box in each panel. Points are colored by brain size residual following the scale at the bottom.

Interestingly, there is no overlap between the bulk motor cortex OCRs and PV+ interneuron OCRs with predicted activity that is significantly associated with brain size residual. In fact, no house mouse OCR ortholog from either set is within 5Mb of a house mouse OCR ortholog from the other set. We also investigated liver OCRs associated with brain size residual and found that none of these OCRs overlapped the associated motor cortex OCRs (**Supplementary Text**) (*26*). This highlights the complementary information provided by using TACIT OCRs from different tissues as well as from using both TACIT and Cell-TACIT.

#### Cell-TACIT and TACIT identify PV+ interneuron and motor cortex open chromatin regions in loci associated with the evolution of social living

One challenge of using TACIT and Cell-TACIT is that tens to hundreds of thousands of OCRs are tested, which requires correcting for large numbers of hypotheses. This is necessary for applying TACIT to phenotypes like brain size for which there is no strict subset of the genome that is known to be involved in the phenotype. In contrast, when such a subset is known, we can increase power by restricting OCRs to those in that subset. We used this targeted approach to examine relationships between solitary and group living lifestyles and predicted PV+ OCR activity within the 1,661,222bp Williams-Beuren Syndrome (WBS) deletion region (**Fig. 6A**), where haploinsufficiency causes increased sociability, intellectual disability, and enhanced verbal fluency in human patients (*69*). Although the WBS locus has not been linked to PV+ interneurons specifically, PV+ interneurons are well-known for their involvement in social behaviors and neuropsychiatric disorders with social components such as autism spectrum disorder (ASD) and schizophrenia (*70*). Molecular evidence for PV+ interneuron involvement suggests associated transcriptional changes. For example, *PVALB* was the most strongly downregulated transcript in ASD brain tissue compared to healthy controls and in animal models of monogenetic neurodevelopmental syndromic disorders (*71, 72*), and single-nucleus RNA-seq from schizophrenia brain tissue revealed more abnormal gene expression in PV+ interneurons than in any other neuronal cell type (*73, 74*). Direct expression manipulation of psychiatric genes in PV+ interneurons was shown to induce social deficits in mice, whereas similar manipulations in other neuron cell types had different effects (*75*).

**Figure 6:**
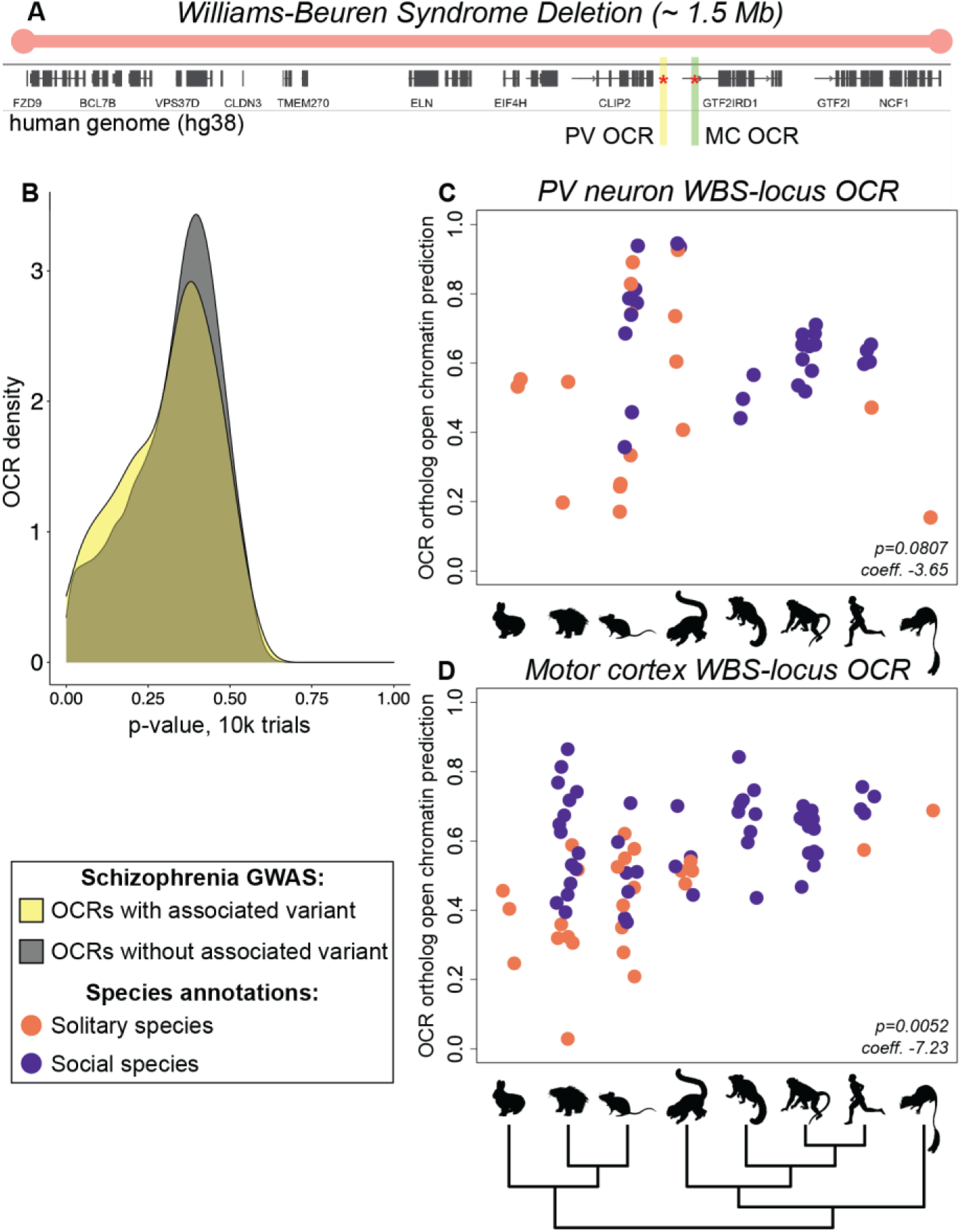
Associations between predicted PV+ interneuron and motor cortex OCR ortholog open chromatin and solitary living. (**A**) A visualization of the human WBS deletion region. The locations of the PV+ interneuron and motor cortex OCRs (highlighted in panels (**C**) and (**D**)) near the gene *GTF2IRD1* are shown in yellow and green, respectively. (**B**) shows the difference in p-value distributions for association between solitary living and predicted open chromatin of PV+ interneuron OCRs whose human ortholog overlaps schizophrenia GWAS SNPs versus all other PV+ interneuron OCRs with a human ortholog. (**C**) highlights the marginal negative association between predicted PV+ interneuron open chromatin and solitary living of orthologs of a PV+ interneuron OCR near *GTF2IRD1*, chr5:134485808-134486308 (mm10). (**D**) highlights the negative association between predicted motor cortex open chromatin and solitary living of orthologs of a motor cortex OCR near *GTF2IRD1*, chr3:42408296-42408946 (rheMac8). For panels (**C-D**), each point represents one ortholog; they are grouped along the x-axis of each panel by clade as shown by the tree below. The clades and example species are listed in Table S10. Points are colored to indicate solitary versus social living following the key at the lower left.

The Mesozoic ancestors of today’s mammals were likely primarily solitary-living, defined by separate foraging and home ranges for females (*76*). Following the End-Cretaceous Mass Extinction, many extant lineages in disparate clades evolved towards social living strategies, including group living and breeding pair monogamy (*76*). Given the impact of PV+ neuron gene expression on social behaviors, we hypothesized that there might be PV+ OCR evolution associated with social structure transitions in mammals.

Before investigating our results, we evaluated whether Cell-TACIT was producing reliable results by comparing results from Cell-TACIT run genome-wide on PV+ OCR orthologs to locations of human genome-wide association study (GWAS) hits for schizophrenia, a disorder associated with solitariness.

Specifically, we divided PV+ OCRs into two groups: those that overlapped a schizophrenia-associated variant and those that did not (*77*). We determined the strength of association of all OCRs with solitary living in mammals. The set of PV+ interneuron OCRs with schizophrenia-associated variants had a shifted phyloglm p-value distribution for association with solitary living compared to the p-value distribution for other PV+ interneuron OCRs (one-sided Wilcoxon rank-sum p = 0.035) (**Fig. 6B**).

When applying Cell-TACIT to only the WBS locus, we identified a mouse OCR (out of two OCRs in this locus) 29kb upstream of *GTF2IRD1* (human ortholog is 36kb upstream) that was marginally associated with non-solitary living (p = 0.08) (**Fig. 6C**) and associated with group living (p = 0.02). To evaluate whether this association was limited to PV+ interneurons, we also evaluated the relationship between predicted bulk motor cortex open chromatin and solitary as well as group living. For solitariness, we found one significantly negatively associated OCR (p = 0.005) (**Fig. 6D**). This OCR is in an intron of *GTF2IRD1* that is about 26kb from its nearest TSS but does not overlap the OCR identified for PV+ interneurons. For group living, we found two significantly associated OCRs, one of which is negatively associated (p = 0.04) and the other of which is positively associated (p = 0.008) and is the same OCR we found for solitariness. Of the 27 protein-coding genes in the WBS locus, *GTF2IRD1* is one of only two genes, where the other gene is its neighbor (*GTF2I*), with structural variants associated with the extreme sociability in dogs that makes them easier to domesticate than wolves (*78*). We additionally evaluated the relationship between predicted liver open chromatin and solitary as well as group living but did not obtain any statistically significant relationships after multiple hypothesis correction.

### TACIT and Cell-TACIT identify open chromatin regions associated with the evolution of vocal learning

We applied TACIT and Cell-TACIT to vocal learning, the ability to modify vocal output as a result of social experience, which has convergently evolved across mammals and been associated with convergent patterns of gene expression in the motor cortex (*2, 79, 80*). We identified 42 OCRs displaying convergent patterns of predicted open chromatin after false discovery rate correction (α=0.05) for motor cortex tissue and 14 for PV+ interneurons, which are described in more depth in our other work . Notably, these vocal learning-associated OCRs showed some concordance with results obtained using complementary methods for detecting convergent evolution. One of the motor cortex OCRs lies 88kb from *Vip*, whose expression in the motor cortex has been associated with vocal learning (*2*). Another OCR is 715kb from *TSHZ3*, whose amino acid sequence also showed convergent evolution associated with vocal learning behavior (p < 0.0001, rank 3) (*81*). *TSHZ3* is involved in the formation of cortico-striatal circuits, which play a central role in vocal learning behavior in mammals and birds, and its disruption in the human population is associated with a form of autism that includes delayed or disrupted speech acquisition (*80, 82*).

## DISCUSSION

We present TACIT and Cell-TACIT, new methods for associating genotypes to phenotypes based on machine learning predictions of tissue- or cell type-specific open chromatin. Our approach overcomes the limitations of nucleotide-level conservation-based approaches, which cannot completely account for the conservation of enhancer function in the presence of low sequence conservation and cannot capture the tissue- and cell type-specificity of enhancer activity (*25*), because our machine learning models learn the conserved regulatory code underlying enhancer activity in our tissue or cell type of interest. We provide a community resource of annotated predicted open chromatin for more than 400,000 OCRs from four tissues and cell types across 222 mammalian species.

We applied TACIT and Cell-TACIT to identify tissue- and cell type-specific OCRs whose predicted open chromatin status across species is associated with brain size residual, solitary living, group living, and vocal learning, including OCRs near genes that were previously implicated in these phenotypes. Specifically, we identified motor cortex and PV+ interneuron OCRs associated with brain size residual that are near genes whose mutations are associated with microcephaly and macrocephaly, as well as motor cortex OCRs with a strong brain size residual association in Cetaceans, which provide candidate mechanisms for the evolution of brain size beyond the previously identified human-specific deletion (*10*). In addition, the WBS deletion region OCRs with the strongest evolution of solitary and group living association are near a critical gene for WBS presentation as well as canine social behavior (*78*). Genome-wide, the associations of PV+ interneuron OCRs with group and solitary living are correlated with whether the OCR overlaps a GWAS hit for schizophrenia, which suggests that OCRs involved in the evolution of traits may also be involved in disorders associated with those traits, a result further supported by our other work (*38*). To be confident that the OCRs we identified have enhancer activity that differs between species, we would need to use reporter assays to test the OCR orthologs’ enhancer activity in multiple species. In addition, to thoroughly demonstrate that these OCRs regulate the nearby genes associated with the phenotypes, we would need to do experiments like CRISPR followed by RNA-qPCR to knock out the OCR and show that the knock-out causes a change in the expression of the nearby gene. Furthermore, considering genes with TSSs within 1Mb may limit our ability to identify real gene-OCR relationships (*83*), but, as data measuring three-dimensional genome interactions becomes available at higher resolution and in additional species, tissues, and cell types, our ability to link candidate enhancers associated with phenotypes to the genes they likely regulate will improve.

While our previous work used data from three species for model-training (*25*), in this work, we developed a new strategy for negative set construction that allowed us to train accurate models using data from only two species. Our success in doing this enabled us to train models that accurately predict whether sequence differences across species in PV+ interneuron OCR orthologs are associated with PV+ interneuron open chromatin changes, demonstrating that the regulatory code is conserved across Euarchontoglires not only at the bulk tissue level but also in a specific neuronal cell type. We also found that having data from more clades enabled us to identify OCRs associated with phenotypes in additional clades, such as the OCR near *LRIG1* associated with the evolution of brain size residual in the Cetacea order within Laurasiatheria, and provides us with the power to identify OCRs with weaker associations with a phenotype across multiple lineages, such as the OCR near *Sall3* associated with the evolution of brain size residual in both Euarchonta and Laurasiatheria.

Unlike phyloP or PhastCons scores, the broad application of TACIT and Cell-TACT is limited by the availability of high-quality open chromatin data from the same tissue or cell type in multiple species. TACIT and Cell-TACIT require enhancer activity data from at least two species for evaluating machine learning models, and, to limit confounding factors, the data should ideally contain animals at comparable developmental stages, biological replicates from both sexes, and animals that were sacrificed in comparable behavioral states at approximately the same relative time in their circadian cycles.

Additionally, predictions are currently limited to orthologs of experimentally identified candidate enhancers, meaning that we are not able to capture enhancers that are not active in the experimentally assayed species, cell types, developmental stages, or conditions. Furthermore, our approach assumes that the evolution of a phenotype is controlled by the same candidate enhancer across species, but there are likely many phenotypes controlled by genes that are not activated by the same enhancer in every species. We also treat missing or unusable OCR orthologs as missing data, but some of these are likely lost OCRs. Exciting extensions to our approach include training models to accurately predict whether sequence differences cause changes in candidate enhancer activity genome-wide, jointly modeling cross-species predicted enhancer activity of enhancers near the same gene, and using genome quality and the predicted open chromatin of OCRs in closely related species to determine when a lack of a usable OCR ortholog should be treated as a negative. Finally, our approach assumes that the regulatory code in our tissue or cell type of interest is conserved across the species we are testing, an assumption that may be violated in some tissues and cell types. For example, this may explain the sub-optimal performance of our retina CNNs trained on mouse sequences in predicting Euarchonta-specific open and closed chromatin (*84, 85*).

With the Zoonomia Cactus alignment of over two hundred mammalian genomes and the wealth of publicly available enhancer activity data from matching tissues and cell types in human, house mouse, and some other species, TACIT and Cell-TACIT can currently be applied to identify candidate enhancers associated with the evolution of many mammalian phenotypes. Because TACIT and Cell-TACIT require enhancer activity data from tissues or cell types of interest in only a few species, they can be used to identify losses of enhancer activity associated with changes in a phenotype in challenging-to-study species for which we have genomes but cannot collect tissue samples. In addition, while we trained our models for TACIT using open chromatin, TACIT can also be applied using other assays of enhancer activity, such as H3K27ac and EP300 ChIP-seq (*27*). Candidate enhancers associated with the evolution of phenotypes near genes involved in diseases related to those phenotypes may provide insights into disease mechanisms. We anticipate that, as more genomes and regulatory genomics data become available, TACIT and Cell-TACIT will provide insights into the regulatory mechanisms governing a wide range of phenotypes.

## Acknowledgements

We would like to thank D. Genereux, D. Levesque, K. Lindblad-Toh, and the members of the Pfenning Lab for useful discussions and suggestions. We would also like to thank P. Sullivan for curating the brain size residual annotations, A. Hindle for providing us with annotations of which mammals spend time underground, and M. Chikina, A. Kowalczyk, and E. Saputra for providing us early access to code for phylogenetic permulations for phylogenetic generalized least squares. This work used the Extreme Science and Engineering Discovery Environment (XSEDE), through the Pittsburgh Supercomputing Center Bridges and Bridges-2 Compute Clusters, which was supported by National Science Foundation grant number TG-BIO200055. Portions of this research were conducted on Lehigh University’s Research Computing infrastructure, which is partially supported by NSF Award 2019035.

## Author contributions

I.M.K., A.J.L., and D.E.S. are listed as co-first authors in last name-alphabetical order because they contributed equally to the manuscript. Conceptualization was done by I.M.K. and A.R.P. Data curation was done by I.M.K. with assistance from C.S., B.N.P., A.J.L., W.K.M., and K.F. Formal analysis was done by I.M.K., D.E.S., and A.J.L. with assistance from C.S. and B.N.P. Funding acquisition was done by A.R.P. with assistance from A.J.L. and B.N.P. Investigation was done by I.M.K., A.J.L., D.E.S., and C.S. with assistance from M.E.W., B.N.P., K.P., A.R.B. and A.R.P. Methodology was developed by I.M.K., A.J.L., and D.E.S. with assistance from A.R.P. Supervision was done by I.M.K. and A.R.P. with assistance from A.J.L.,W.K.M., and M.E.W. Software was implemented by D.E.S, I.M.K., A.J.L., and C.S. with assistance from M.E.W., W.K.M., X.Z., and K.F. Visualization was done by I.M.K. and D.E.S. with assistance from C.S., A.J.L., and A.R.P. Manuscript preparation was done by I.M.K., D.E.S., and A.J.L. with assistance from A.R.P. and C.S. Manuscript review and editing was done by all authors.

## Data and materials availability

Publicly available ATAC-seq data was obtained from Gene Expression Omnibus accessions GSE161374, GSE146897, and GSE137311; China National GeneBank accession CNP0000198; and ArrayExpress accession E-MTAB-2633. Unpublished ATAC-seq data generated by the Pfenning Lab can be found at GSE159815 with token gzkbqwyobvmvfsd and will be released prior to publication. The tree used for the phenotype association pipeline can be obtained by contacting the Zoonomia Consortium and will be released prior to publication. Publicly available genomes and annotations were downloaded from NCBI Assembly and the UCSC Genome Browser. Publicly available Hi-C data was downloaded from http://hugin2.genetics.unc.edu/Project/hugin/. Motif discovery results and machine learning models can be found at http://daphne.compbio.cs.cmu.edu/files/ikaplow/TACITSupplement/. Machine learning model predictions can be obtained from authors upon request and will be released prior to publication. Code also used in our previous work (*25*) can be found at https://github.com/pfenninglab/OCROrthologPrediction, new code for this work can be found at https://github.com/pfenninglab/TACIT, and additional code can be obtained from authors upon request.

## ZOONOMIA CONSORITUM MEMBERS

Gregory Andrews^1^, Joel C. Armstrong^2^, Matteo Bianchi^3^, Bruce W. Birren^4^, Kevin R Bredemeyer^5^, Ana M Breit^6^, Matthew J Christmas^3^, Joana Damas^7^, Mark Diekhans^2^, Michael X. Dong^3^, Eduardo Eizirik^8^, Kaili Fan^1^, Cornelia Fanter^9^, Nicole M. Foley^5^, Karin Forsberg-Nilsson^10^, Carlos J. Garcia^11^, John Gatesy^12^, Steven Gazal^13^, Diane P. Genereux^4^, Daniel Goodman^14^, Linda Goodman^15^, Jenna Grimshaw^11^, Michaela K. Halsey^11^, Andrew J Harris^5^, Glenn Hickey^16^, Michael Hiller^17^, Allyson Hindle^9^, Robert M. Hubley^18^, Graham Hughes^19^, Jeremy Johnson^4^, David Juan^20^, Irene M. Kaplow^21,22^, Elinor K. Karlsson^1,4^, Kathleen C. Keough^23,24^, Bogdan Kirilenko^17^, Jennifer M. Korstian^11^, Sergey V. Kozyrev^3^, Alyssa J. Lawler^25^, Colleen Lawless^19^, Danielle L. Levesque^6^, Harris A. Lewin ^7,26,27^, Xue Li^1,4^, Abigail Lind^23,24^, Kerstin Lindblad-Toh^3,4^, Voichita D. Marinescu^3^, Tomas Marques-Bonet^20,28,29,30^, Victor C Mason^31^, Jennifer R. S. Meadows^3^, Jill Moore^1^, Diana D. Moreno-Santillan^11^, Kathleen M. Morrill^1,4^, Gerard Muntané^20^, William J Murphy^5^, Arcadi Navarro^20,32,33,34^, Martin Nweeia^35,36,37,38^, Austin Osmanski^11^, Benedict Paten^2^, Nicole S. Paulat^11^, Eric Pederson^3^, Andreas R. Pfenning^21,22^, BaDoi N. Phan^21^, Katherine S. Pollard^23,24,39^, Kavya Prasad^21^, Henry Pratt^1^, David A. Ray^11^, Jeb Rosen^18^, Irina Ruf ^40^, Louise Ryan^19^, Oliver A. Ryder^41,42^, Daniel Schäffer^21^, Aitor Serres^20^, Beth Shapiro^43,44,^ Arian F. A. Smit^18^, Mark Springer^45^, Chaitanya Srinivasan^21^, Cynthia Steiner^46^, Jessica M. Storer^18^, Patrick F. Sullivan^47,48^, Kevin A. M. Sullivan^10^, Elisabeth Sundström^3^, Megan A Supple^44^, Ross Swofford^4^, Joy-El Talbot^49^, Emma Teeling^19^, Jason Turner-Maier^4^, Alejandro Valenzuela^20^, Franziska Wagner^40^, Ola Wallerman^3^, Chao Wang^3^, Juehan Wang^13^, Zhiping Weng^1^, Aryn P. Wilder^41^, Morgan E. Wirthlin^21,22^, Shuyang Yao^48^, Xiaomeng Zhang^21^

^1^Program in Bioinformatics and Integrative Biology, University of Massachusetts Medical School, Worcester, MA 01605, USA

^2^Genomics Institute, UC Santa Cruz, 1156 High Street, Santa Cruz, CA 95064, USA

^3^Science for Life Laboratory, Department of Medical Biochemistry and Microbiology, Uppsala University, Uppsala, 751 32, Sweden

^4^Broad Institute of MIT and Harvard, Cambridge MA 02139, USA

^5^Veterinary Integrative Biosciences, Texas A&M University, College Station, TX 77843, USA

^6^School of Biology and Ecology, University of Maine, Orono, Maine 04469, USA

^7^The Genome Center, University of California Davis, Davis, CA 95616, USA

^8^School of Health and Life Sciences, Pontifical Catholic University of Rio Grande do Sul, Porto Alegre, 90619-900, Brazil

^9^School of Life Sciences, University of Nevada Las Vegas, Las Vegas, NV 89154, USA

^10^Department of Immunology, Genetics and Pathology, Science for Life Laboratory, Uppsala University, Uppsala, 751 85, Sweden

^11^Department of Biological Sciences, Texas Tech University, Lubbock, TX 79409, USA

^12^Division of Vertebrate Zoology, American Museum of Natural History, New York, NY 10024, USA

^13^Keck School of Medicine, University of Southern California, Los Angeles, CA 90033, USA

^14^University of California San Francisco, San Francisco, CA 94143 USA

^15^Fauna Bio Inc., Emeryville, CA 94608, USA

^16^Baskin School of Engineering, University of California Santa Cruz, Santa Cruz, CA 95064, USA

^17^Max Planck Institute of Molecular Cell Biology and Genetics, 01307, Dresden, Germany

^18^Institute for Systems Biology, Seattle, WA 98109, USA

^19^School of Biology and Environmental Science, University College Dublin, Belfield, Dublin 4, Ireland

^20^Institute of Evolutionary Biology (UPF-CSIC), Department of Experimental and Health Sciences, Universitat Pompeu Fabra, Barcelona, 08003, Spain

^21^Department of Computational Biology, School of Computer Science, Carnegie Mellon University, Pittsburgh, PA 15213, USA

^22^Neuroscience Institute, Carnegie Mellon University, Pittsburgh, PA 15213, USA

^23^Gladstone Institutes, San Francisco, CA 94158, USA

^24^Department of Epidemiology & Biostatistics, University of California, San Francisco, CA 94158, USA

^25^Department of Biology, Carnegie Mellon University, Pittsburgh, PA 15213, USA

^26^Department of Evolution and Ecology, University of California, Davis, CA 95616, USA

^27^John Muir Institute for the Environment, University of California, Davis, CA 95616, USA

^28^Catalan Institution of Research and Advanced Studies (ICREA), 08010, Barcelona, Spain

^29^CNAG-CRG, Centre for Genomic Regulation, Barcelona Institute of Science and Technology (BIST), 08036, Barcelona, Spain

^30^Institut Català de Paleontologia Miquel Crusafont, Universitat Autònoma de Barcelona, 08193, Cerdanyola del Vallès, Barcelona, Spain

^31^Institute of Cell Biology, University of Bern, 3012 Bern, Switzerland

^32^Catalan Institution of Research and Advanced Studies (ICREA), 08010, Barcelona, Spain

^33^CRG, Centre for Genomic Regulation, Barcelona Institute of Science and Technology (BIST), 08036, Barcelona, Spain

^34^BarcelonaBeta Brain Research Center, Pasqual Maragall Foundation, Barcelona, 08005 Spain

^35^Narwhal Genome Initiative, Department of Restorative Dentistry and Biomaterials Sciences, Harvard School of Dental Medicine, Boston, MA 02115, USA

^36^Department of Comprehensive Care, School of Dental Medicine, Case Western Reserve University, Cleveland, OH 44106, USA

^37^Department of Vertebrate Zoology, Smithsonian Institution, Washington, DC 20002, USA

^38^Department of Vertebrate Zoology, Canadian Museum of Nature, Ottawa, Ontario K2P 2R1, Canada

^39^Chan Zuckerberg Biohub, San Francisco, CA 94158, USA

^40^Division of Messel Research and Mammalogy, Senckenberg Research Institute and Natural History Museum Frankfurt, 60325 Frankfurt am Main, Germany

^41^Conservation Genetics, San Diego Zoo Wildlife Alliance, Escondido, CA 92027, USA

^42^Department of Evolution, Behavior and Ecology, Division of Biology, University of California, San Diego, La Jolla, CA 92039 USA

^43^Howard Hughes Medical Institute, University of California Santa Cruz, Santa Cruz, CA 95064, USA

^44^Department of Ecology and Evolutionary Biology, University of California Santa Cruz, Santa Cruz, CA 95064, USA

^45^Department of Evolution, Ecology and Organismal Biology, University of California, Riverside, CA 92521, USA

^46^Conservation Science Wildlife Health, San Diego Zoo Wildlife Alliance, Escondido CA 92027, USA

^47^Department of Genetics, University of North Carolina Medical School, Chapel Hill, NC 27599, USA

^48^Department of Medical Epidemiology and Biostatistics, Karolinska Institute, Stockholm, Sweden

^49^Iris Data Solutions, LLC, Orono, ME 04473, USA

