## Supplement for "Relating enhancer genetic variation across mammals to complex phenotypes using machine learning"

### Supplementary Materials for Relating enhancer genetic variation across mammals to complex phenotypes using machine learning

#### Materials and Methods

##### *Machine learning model training and evaluation*

<u>Data processing</u>

Bulk ATAC-seq data:

We obtained processed bulk ATAC-seq data from motor cortex, striatum, liver, and retina from our and others' previous work. Specifically, we obtained processed house mouse bulk motor cortex and striatum ATAC-seq data from GSE161374 (*88*). We obtained house mouse bulk liver ATAC-seq data from CNP0000198 (*89*) and GSE159815 (*25*) and used the processed form of these datasets from our previous work (*25*). We obtained rhesus macaque and Norway rat bulk motor cortex (orofacial motor cortex and hand motor cortex), striatum, and liver ATAC-seq from GSE159815 (*90*) and used the processed form of these datasets from our previous work (*25*) with the exception of rhesus macaque striatum, for which we used the the IDR optimal peaks from putamen that intersect IDR optimal peaks from caudate (the difference from our previous work was not also requiring intersection with peaks from nucleus accumbens) obtained with bedtools (*91*) so that it would be more directly comparable to the Norway rat (mostly putamen) (*90*) and Egyptian fruit bat (We also used the putamen IDR optimal peaks that intersected caudate IDR optimal peaks.) (*81*) striatum data. We used the processed form of the Egyptian fruit bat bulk motor cortex (orofacial motor cortex and wing motor cortex) and striatum (putamen and caudate) ATAC-seq data from (*81*). We obtained cow and pig bulk liver ATAC-seq data from E-MTAB-2633 (*92*) and used the processed form of these datasets from our previous work (*25*). We obtained house mouse retina data from GSE146897 (*93*) and human retina data from GSE137311 (*94*) and processed the datasets using ENCODE ATAC-seq pipeline v1.9.1 with default parameters except for "atac.auto_detect_adapter":true, "atac.multimapping":0, and "atac.idr_thresh":0.1, where we mapped the reads to the mm10 and hg38 assemblies, respectively, to obtain IDR "optimal" peaks.

Single-nucleus ATAC-seq data:

We obtained house mouse motor cortex single-nucleus ATAC-seq (snATAC-seq) (*95*) and human motor cortex Single-nucleus Chromatin Accessibility and mRNA Expression Sequencing 2 (SNARE-seq2) ATAC-seq (*96*) data from NEMO. We aligned the house mouse snATAC-seq reads to the mm10 assembly (*97*) using bowtie2 (*98*) with default parameters and used the bam files with human SNARE-seq2 ATAC-seq reads aligned to the hg38 assembly (*99*). Next, we sorted the bam files and created bam indices with samtools (*100*) and then, for the human data, used alignmentSieve (*101*) with settings --minMappingQuality 10, --minFragmentLength 10, and --ignoreDuplicates to remove multi-mappers and especially short reads. We created a new bam index for the human bam file with samtools (*100*) and then used ArchR's createArrowFiles (*102*) with parameters filterTSS = 0 and filterFrags = 0 for house mouse and filterTSS = 3 and filterFrags = 500 for human to convert the bam files into arrow files. After that, we used the arrow files to infer the doublet scores for each cell using ArchR's addDoubletScores with default parameters and then filtered the doublets using ArchR's filterDoublets with setting cutEnrich = 5 (*102*). We filtered the cells that were removed in the work that presented the datasets and clustered the cells according to the cluster labels in that work (*95, 96, 103*). Finally, we merged the cells in each cluster to create a single coverage file for each cluster with ArchR's addGroupCoverages with default parameters and identified reproducible peaks with ArchR's addReproduciblePeakSet with default parameters (*102*).

Constructing positive sets

To obtain OCRs, we intersected the reproducible peaks from each of the motor cortex regions for motor cortex and each of the liver datasets for liver and then, for each species defined OCRs to be the intersected peaks that are likely to be enhancers. Specifically, for each species, we selected one set of reproducible open chromatin peaks to be the “base peaks,” used bedtools v2.29 intersect with the -wa and -u options to intersect it with each of the other reproducible peak sets in series, and then used bedtools v2.29 closestBed with the -t first (we did not use this option for house mouse motor cortex, but we removed all repeated OCRs before running phylolm/pholglm and doing test set evaluations) and -d options to identify the “base peaks" that overlapped at least one peak from each other set, that were over 20kb from the nearest protein-coding TSS (not promoters, we used the nearest protein-coding transcript for house mouse motor cortex), at most 1kb long (not super-enhancers), and non-exonic (*91*). The base peaks for house mouse motor cortex were the irreproducible discovery rate (IDR) (*104, 105*) “optimal set” peaks from bulk motor cortex data we previously generated (*88*), for the house mouse liver were the IDR “optimal set” from the house mouse liver data we previously generated (*90*), for the house mouse PV+ interneurons were the reproducible PV+ interneuron peaks from the mouse snATAC-seq data (*95, 103*), for the house mouse retina were the IDR “optimal set” from the house mouse retina data we processed (*93*), for the rhesus macaque motor cortex were the IDR “optimal set” from the orofacial motor cortex data we previously generated (*90*), for the rhesus macaque liver were the IDR reproducible peaks across self-pseudo replicates from the first rhesus macaque liver replicate from data we previously generated (*90*), for the Norway rat brain were the IDR “optimal set” from the primary motor cortex from data we previously generated (*90*), for the Norway rat liver were the IDR “optimal set” from the second and third Norway rat liver replicates from data we previously generated (*90*), for the Egyptian fruit bat motor cortex were the IDR “optimal set” from the orofacial motor cortex data presented in our other work (*81*), for cow and pig liver were the IDR “optimal set” generated by the FAANG consortium (*92*) and processed in our previous work (*25*), for human PV+ interneurons were the reproducible PV+ interneuron peaks from the human SNARE-seq2 ATAC-seq data (*96, 103*) were the IDR “optimal set” from the human retina data we processed (*94*). To determine the distance from the nearest protein-coding TSS, we used the house mouse GENCODE protein-coding TSSs for house mouse (version M15) (*106*), the union of the RefSeq rheMac8 protein-coding TSSs (*107*) and the human GENCODE (version 27) protein-coding TSSs mapped to rheMac8 using liftOver (*108*) for rhesus macaque, the union of the RefSeq rn6 protein-coding TSSs (*107*) and the house mouse GENCODE (version M15) protein-coding TSSs mapped to rn6 using liftOver (*108*) for Norway rat, the union of RefSeq mRouAeg1 TSSs (no clear annotation for protein-coding TSSs) and the human GENCODE (version 27) protein-coding TSSs mapped to mRouAeg1 with liftOver (*108*) (We used liftOver to mRouAge1 instead of halLiftover (*13*) to the Egyptian fruit bat alignment in the Zoonomia Cactus (*12*) followed by liftOver to mRouAeg1 because mRouAeg1 has substantially fewer gaps than other bat assemblies (*109*).) for Egyptian fruit bat, the union of the RefSeq Btau_5.0.1 TSSs (*107*) and the human GENCODE protein-coding TSSs mapped to Btau_5.0.1 with halLiftover (*13*) on the version 1 Zoonomia Cactus alignment (*12*) for cow, the union of the susScr11 TSSs mapped to susScr3 with liftOver (*108*) and the human GENCODE (version 27) protein-coding TSSs mapped to susScr3 with halLiftover (*13*) on the version 1 Zoonomia Cactus alignment (*12*) for pig, and the human GENCODE (version 27) protein-coding TSSs for human (*106*). To identify non-exonic peaks, we used bedtools (*91*) subtract with option -A to identify peaks that did not overlap protein-coding exons, where we obtained house mouse protein-coding exons from GENCODE (version M15) (*106*), rhesus macaque protein-coding exons from RefSeq for rheMac8 (*107*), Norway rat protein-coding exons from RefSeq for rn6 (*107*), Egyptian fruit bat protein-coding exons from RefSeq for mRouAeg1, cow protein-coding exons from RefSeq for Btau_5.0.1 (*107*), pig protein-coding exons from RefSeq for susScr11 (*107*) and mapped to susScr3 with liftOver (*108*), and human protein-coding exons from GENCODE (version 27) (*106*). We defined the peak summit of an OCR to be the peak summit of the corresponding base peak, and we constructed positive examples by taking OCR peaks summits +/- 250bp and their reverse complements. We centered peaks on their summits because we and others have previously shown that there is a concentration of TF motifs at peak summits (*14*, *110*).

##### Constructing negative sets

Constructing negative set with non-OCR orthologs of OCRs for tissues with data from more than two species:

When we had data from at least three species, we used non-OCR orthologs of OCRs as our negative set because CNNs trained on this negative set have previously been shown to have high lineage-specific OCR accuracy, tissue-specific OCR accuracy, and phylogeny-matching correlations (*25*). To ensure that we would have enough negatives for training the bulk open chromatin models, for each species, tissue combination with multiple datasets, we created a less conservative set of OCRs, which we called "loose OCRs." The loose OCRs are the base peaks that are non-exonic, at least 20kb from a TSS, and at most 1kb long (same criteria used in constructing the positive set) and, if applicable, intersect at least 1 peak from the pooled reads across replicates from the other dataset that we used for the species, tissue combination. We obtained these loose OCRs using bedtools v2.29 (*91*) and defined the peak summit of a loose OCR to be the peak summit of the corresponding base peak. We identified orthologs of our loose OCRs in each other species with open chromatin data from the same tissue using halLiftover (*13*) with the Zoonomia version 2 Cactus alignment (*12*) followed by HALPER (*14*) with parameters -max_frac 2.0, -min_len 50, and -protect_dist 5. We then used bedtools subtract with the -A option (*91*) to remove the loose OCR orthologs in each species that overlapped peaks from the pooled reads across replicates from any of the datasets from the same tissue in that species with the exception of non-house mouse bulk motor cortex loose OCR orthologs in mouse, for which we used the same approach to remove loose OCR orthologs overlapping peaks from pooled replicates from the separate age group processing described in our previous work (*25*). To construct negatives from the non-OCR orthologs of loose OCRs, we used the sequence underlying the ortholog of the base peak summit +/- 250bp and that sequence's reverse complement. Our negative:positive training set ratios were approximately 0.695:1 for the motor cortex CNN trained on only house mouse sequences, 1.09:1 for the multi-species motor cortex CNN, and 0.728:1 for the multi-species liver CNN.

Constructing negative set with G/C-content-matched regions and OCRs from other cell types for models for cell types with data from only two species:

For tissues and cell types with data from only two species, non-OCR orthologs of OCRs did not seem to provide a sufficient quantity of negatives for accurate model training, so we constructed negative sets that were a combination of 1) a large number of closed chromatin regions with similar G/C-content to the positive set and 2) genomic regions with accessible chromatin in other motor cortex neuron cell types but not the cell type used to train the models. For PV+ interneurons, we obtained GC-, repeat-matched sequences using the genNullSeqs() function from the gkmSVM R package, version 0.81.0, with default settings except for increasing xfold to 10 and nMaxTrials to 100 to identify a large number of candidate sequences and decreasing length_match_tol to 0.0 to require output sequences of exactly 500 bp (*111*). We used masked genomes from R Bioconductor packages BSgenome.Mmusculus.UCSC.mm10.masked version 1.3.99 for house mouse and BSgenome.Hsapiens.UCSC.hg38.masked version 1.3.993 for human to avoid tandem repeat regions, intra-contig ambiguities, and assembly gaps (*112*). We filtered the results to remove possible promoter regions within 20 kb of a TSS using the same process that we used for constructing the positive set. For the negative examples from accessible chromatin in other motor cortex neuron cell types, we used the process that we used for constructing the positive sets to obtain 500 bp regions surrounding non-promoter (> 20kb from the nearest TSS), non-super-enhancer (≤ 1kb long) open chromatin peak summits from each of the following neuron cell types: Sst, Vip, Lamp5, Chodl, Sncg, NP, L23, L5.IT, L5.PT, L6.CT, and L6.IT. Finally, to remove any potential positive contamination from the combined negative set, we removed G/C, repeat-matched negatives with direct overlap with OCRs in the cell type used in model training (including non-reproducible OCRs) as well as OCRs from other cell types that were within 5 kb of OCRs (including non-reproducible OCRs) in the cell type used for model training. Additionally, we removed regions from the combined negative set that intersected any ATAC-seq peaks from pooled replicates from isolated PV+ interneuron populations from our previous work (*60*). We did these operations using bedtools v2.29 (*91*). The mouse-only PV+ interneuron CNN training set had negative:positive ratio 4.77:1, and the multi-species PV+ interneuron CNN training set had negative:positive ratio 4.91:1.

For the retina models, we generated four times the number of positive sequences that are G/C-matched and not overlapping mouse retina peaks. We did this by first building a background sequence repository from the positive sequences using BiasAway's biasaway_background_construction function (*113, 114*). Next, we used biasaway (*113, 114*) to generate negative sequences with a matching nucleotide distribution along a sliding window with the following settings: --nfold 4 --deviation 2.6 --step 50 --seed 1 --winlen 100. Then, we used bedtools to remove the negative sequences that overlapped mouse retina IDR "optimal set" peaks. Finally, we added the positives from the mouse-only motor cortex models that did not overlap any retina IDR "optimal set" peaks. The mouse-only retina CNN training set had negative:positive ratio 4:1, and the multi-species PV+ interneuron CNN training set had negative:positive ratio 4:1.

Constructing training, validation, and test sets

We constructed training, validation, and test sets using an approach designed to prevent overlap between them. For the model training and evaluation using house mouse sequences, our training set consisted of regions on mm10 chromosomes 3-7, 10-19, and X; our validation set that we used for developing our positive and negative set definitions, early-stopping, and hyper-parameter tuning consisted of regions on mm10 chromosomes 8-9; and our test set consisted of regions on mm10 chromosomes 1-2. We conducted all "test set" model performance evaluations on house mouse genomic regions, including those on types of regions not used in model training, on only regions from mm10 chromosomes 1-2. For the model training and evaluations using sequences from non-house mouse species, we divided sequences into training, validation, and test sets based whether their house mouse orthologs found by mapping them to mm10 using halLiftover (*13*) with the Zoonomia version 2 Cactus alignment (*12*) followed by HALPER (*14*) with parameters -max_frac 2.0, -min_len 50, and -protect_dist 5 mapped to house mouse training, validation, or test set chromosomes. We excluded OCRs with no usable house mouse orthologs because they may have house mouse orthologs not detected due to imperfect alignments.

Training CNNs

We used CNNs (*29*) because they can learn non-linear relationships between sequence patterns, they do not require an explicit featurization of the data, and we have previously shown that they work well for predicting open chromatin status of OCR orthologs (*25*). Our inputs were one-hot-encoded DNA sequences (*115*) (four by 500 matrices), and our outputs were the probability that the input sequence is an OCR in the tissue or cell type for which we trained the CNN. We tried training each CNN using the architecture for the most relevant task from (*25*) and then tuned the learning rate, number of convolutional filters, dropout applied to the convolutional layers, number of fully connected layer units per fully connected layer, and number of fully connected layers by comparing the validation set performances. For the CNNs for models with data from only two species, we additionally tuned the convolutional filter width, momentum, regularization applied to convolutional layers, dropout applied to fully connected layers, and regularization applied to fully connected layers. We implemented and trained all CNNs using Keras (*116*), with version 1.2.2 with the Theano backend (*117*) for CNNs for tissues with data from at least three species, version 2.2.0 with the Theano backend (*117*) for CNNs for PV+ interneurons, and version 2.3.0-tf with the TensorFlow version 2.2.0 (*118*) backend for retina, and we evaluated CNNs using Scikit-learn (*119*) and PRROC (*120*). Each CNN had a series of convolutional layers, followed by a max-pooling layer, followed by one or more fully connected layers. Unless otherwise specified, we initialized the CNN weights with Keras's He normal initializer (*116*, *121*) with the exception of the final fully connected layer, which we initialized using Keras's Glorot normal initializer (*116*, *122*), and used a rectified linear unit as the activation for each layer with the exception of the final fully connected layer, which went into a sigmoid. For motor cortex, liver, and PV+ interneurons, we set each class's weight to be the fraction of examples from the other class in the training set. For retina, the positives were weighted by the product of half of the total and the reciprocal of the number of positives, and the negatives were weighted by the product of half of the total and the reciprocal of the number of negatives.

All CNNs for tissues with data from at least three species had 5 convolutional layers with filters of width 7, stride 1, and L2 regularization 0.000001 in each, which were followed by a max-pooling layer with width and stride 26. We initialized the weights to be those from a pre-trained CNN with the same hyper-parameters and the larger of the positive and negative sets randomly down-sampled to be the size of the other set. We trained the CNNs using stochastic gradient descent with learning rate 0.0005, Nesterov momentum 0.99, and batch size 100 until there were 3 consecutive epochs with no improvement at recall at eighty percent precision (or specificity at eighty percent NPV if there were more positives than negatives) on the validation set.

Our final architecture for the house mouse-only motor cortex CNN was 300 filters per convolutional layer with convolutional layer dropout 0.1, followed by the max-pooling layer, followed by two fully connected layers with 500 units each, followed by a fully connected layer that went into a sigmoid (**Supplementary Website**). Our final architecture for the multi-species motor cortex CNN was 600 filters per convolutional layer with convolutional layer dropout 0.2, followed by the max-pooling layer, followed by one fully connected layer with 300 units each, followed by a fully-connected layer that went into a sigmoid (**Supplementary Website**). Our final architecture for the multi-species liver CNN was 450 filters per convolutional layer with convolutional layer dropout 0.2, followed by the max-pooling layer, followed by one fully connected layer with 300 units each, followed by a fully-connected layer that went into a sigmoid (**Supplementary Website**).

Our hyper-parameters for the house mouse-only and multi-species PV+ interneuron CNNs were the same except for the number of convolutional filters per convolutional layer, the L2 regularization for each layer, and the dropout for each layer. Our final architecture for both PV+ interneuron CNNs was 3 convolutional layers with filters of size 7 and stride 1, followed by a max-pooling layer with size 26 and stride 26, followed by a fully connected layer with 300 units, followed by a fully connected layer without any regularization or dropout that went into a sigmoid. For the house mouse-only CNN, each convolutional layer had 300 filters; the L2 regularization for each convolutional and the fully connected layer was 0.0001; and the dropout was 0.3 for the first convolutional layer, 0.25 for the second convolutional layer, 0.2 for the third convolutional layer, and 0.1 for the first fully connected layer (**Supplementary Website**). For the multi-species CNN, each convolutional layer had 200 filters, the L2 regularization for each convolutional and the fully connected layer was 0.0002, and the dropout was 0.3 for each convolutional layer and for the fully connected layer (**Supplementary Website**). We trained the CNNs using stochastic gradient descent with a cyclic learning rate and momentum (*123*), where the learning rate range was 0.00001 to 0.01, the momentum range was 0.90 to 0.99, and the number of cycles was 2.5; we used batch size 128 and trained CNNs for 60 epochs.

Our hyper-parameters for the mouse-only and multi-species retina models were the same. Our final architecture was 3 convolutional layers with 256, 128, and 64 filters of size 11 and stride 1; followed by a max-pooling layer with size 26 and stride 26; followed by a fully connected layer with 300 units; followed by a fully connected layer without any dropout that went into a sigmoid. The L2 regularization for each convolutional and each fully connected layer was 0.00000001. The dropout was 0.4 for the first convolutional layer, 0.3 for the second convolutional layer, 0.2 for the third convolutional layer, and 0.4 for the first fully connected layer (**Supplementary Website**). We trained the CNNs using stochastic gradient descent with a cyclic learning rate and momentum (*123*), where the learning rate range was 0.0001 to 0.1, the momentum range was 0.85 to 0.99, and the number of cycles was 2.5; we used batch size 128 and trained CNNs for 35 epochs.

Evaluating CNNs

Evaluating CNNs' lineage-specific OCR accuracy:

We defined a clade-specific OCR in a tissue as an OCR whose ortholog is open in that tissue in every species within a clade for which we have data and closed (does not overlap any peaks from the pooled reads across replicates) in every other species for which we have data, and we defined a species-specific OCR as an OCR whose ortholog is open a species for which we have data and is closed (does not overlap any peaks from the pooled reads across replicates) in the most closely related species for which we have data. For some combinations of tissues/cell types and clades, we modified these definitions slightly so that there would be at least one hundred examples (fifty sequences and each sequence's reverse complement) going into each test set evaluation. More specifically, we defined Glires-specific OCRs to be house mouse OCRs in the tissue/cell type whose Norway rat ortholog overlaps an open chromatin peak from the pooled reads across replicates in that tissue/cell type if Norway rat data is available and whose orthologs in all other species do not overlap any peaks from the pooled reads across replicates in that tissue/cell type. We defined Glires-specific non-OCRs to be house mouse orthologs of rhesus macaque (motor cortex and liver)/human (PV+ interneuron and retina) OCRs whose orthologs in any other non-Glires species with available data overlap an open chromatin peak from the pooled reads across replicates and whose orthologs in all Glires species do not overlap any peaks from the pooled reads across replicates. For motor cortex, PV+ interneurons, and retina, we defined Euarchonta-specific OCRs to be rhesus macaque (motor cortex)/human (PV+ interneurons and retina) OCRs whose orthologs in Glires species with available data do not overlap any open chromatin peaks from the pooled reads across replicates. For liver, we added the additional restriction that the orthologs in non-Euarchontoglires species with available data not overlap any open chromatin peaks from the pooled reads across replicates. For motor cortex, PV+ interneurons, and retina, we defined Euarchonta-specific non-OCRs to be rhesus macaque (motor cortex)/human (PV+ interneurons and retina) orthologs of house mouse OCRs that do not overlap any peaks from the pooled reads across replicates and, for motor cortex, whose rat orthologs do overlap a peak from the pooled reads across replicates. For liver, we added the additional restriction that the orthologs in non-Euarchontoglires species with available data overlap open chromatin peaks from the pooled reads across replicates in each of these species. We defined Laurasiatheria-specific OCRs to be

Egyptian fruit bat (motor cortex)/cow (liver OCRs) whose orthologs in each Euarchontoglires species with available data do not overlap any peaks from the pooled reads across replicates and, for liver, whose ortholog in pig overlaps a peak from the pooled reads across replicates. We defined Laurasiatheria-specific non-OCRs to be Egyptian fruit bat (motor cortex)/cow (liver) orthologs of mouse OCRs whose orthologs in Laurasiatheria species with available data do not any peaks from the pooled reads across replicates and whose orthologs in each Euarchontoglires species with available data do overlap peaks from the pooled reads across replicates. When evaluating multi-species CNNs, we combined species-specific OCRs and non-OCRs across species. We obtained these sets of OCRs and non-OCRs using halLiftover (*13*) with the Zoonomia version 2 Cactus alignment (*12*) followed by HALPER (*14*) with parameters -max_frac 2.0, -min_len 50, and -protect_dist 5 and bedtools v2.29 (*91*).

Evaluating CNNs' tissue-specific OCR accuracy:

We evaluated tissue-specific OCR accuracy in two ways. First, we evaluated performance on a very different tissue from the tissue/cell type used in model training, where our positives were OCRs in both tissues (ensures that model is not learning only tissue-specific open chromatin sequence signatures), and our negatives were OCRs in the tissue not used in training that do not overlap any peaks from pooled replicates in the tissue/cell type used in training (ensures that model is learning sequences specific to tissue used in training). The additional tissues we used were liver for motor cortex, PV+ interneuron, and retina CNNs and motor cortex for liver CNNs. Second, we evaluated performance on a similar tissue/cell type to the tissue/cell type used in model training, where our positives were OCRs in both tissues/cell types, and our negatives were OCRs in the tissue/cell type not used in model training that do not overlap any peaks from the pooled replicates from the tissue/cell type used in model training. The additional tissues/cell types we used were striatum for motor cortex CNNs and the union of excitatory neurons (NP, L23, L5.IT, L5.PT, L6.CT, and L6.IT) for PV+ interneuron CNNs; unfortunately, there was no similar tissue to liver or retina with open chromatin data available. We obtained our positives using bedtools v2.29 intersectBed with options -wa and -u, and we obtained our negatives using bedtools v2.29

subtractBed with option -A (*91*). We combined the data from all species for evaluating the multi-species CNNs.

Evaluating if CNNs' predictions had phylogeny-matching correlations:

To evaluate the relationship between OCR ortholog predicted open chromatin status and phylogenetic distance, we identified test set house mouse OCR orthologs in all of the 56 Glires species from Zoonomia, predicted the open chromatin statuses of those orthologs, and investigated the relationships between those predictions and the species' phylogenetic divergences from house mouse. Specifically, we first mapped OCRs across Glires using halLiftover (*13*) with the Zoonomia version 2 Cactus alignment (*12*) followed by HALPER (*14*) with parameters -max_frac 2.0, -min_len 50, and -protect_dist 5. We next constructed inputs for our CNNs, made predictions, and filtered predictions as described in our previous work (*25*). Then, for each CNN, species combination, we computed the mean OCR ortholog open chromatin status prediction and the standard deviation of predictions across all filtered OCR orthologs. After that, we computed the Pearson and Spearman correlations between these means and standard deviations of predictions and the millions of years since divergence from house mouse, which we obtained from TimeTree (*36*). We found that a few species had substantially lower means and standard deviations of predictions than those of species with similar levels of relatedness to house mouse and that many those species's OCR orthologs had a high frequency of N's, making their predictions more likely to be inaccurately negative, so we repeated this process after filtering out all orthologs for which more than five percent of the nucleotides were N's.

Interpreting machine learning models:

To further evaluate whether our motor cortex and liver CNNs had learned sequence patterns related to gene regulation in motor cortex and liver, respectively, we interpreted our CNNs using deepLIFT (*30*) followed by TF-MoDISco (*124*). We first computed the importance of every nucleotide in every example in the validation set using deepLIFT version 0.5.5-theano with the Rescale rule scores from the sequence layer with the target of the final convolutional layer, where our reference was a sequence of N's. We also computed the "hypothetical scores" for each nucleotide at each position for each sequence using an extension to deepLIFT (*124*) with the Rescale rule. We next created motifs by combining the scores and hypothetical scores using TF-MoDISco version 0.5.5.6 with the following settings: seqlet FDR threshold = 0.2; gapped k-mer settings for similarity computation k-mer length = 8, number of gaps = 1, and number of mismatches = 0; and final motif width = 50. We visualized our motifs using the aggregated hypothetical scores of the seqlets supporting each motif and created position frequency matrices by averaging the one-hot-encoded sequences at all of the seqlet coordinates belonging to the motifs. Finally, we compared each motif to known motifs by running TomTom (*125*) on the corresponding position frequency matrix with the CIS-BP version 2.00 *Homo sapiens* motif database (*86*).

To further evaluate whether our retina CNNs had learned sequence patterns related to gene regulation in the retina, we interpreted our CNNs using Deep SHAP (*31*) followed by TF-MoDISco (*124*). We first computed the importance of every nucleotide in 2,000 random examples from the mouse validation set as well as the orthologs of the OCR near *Otx2* that were shown to have accelerated evolution in subterranean mammals (*9*) in subterranean mammal (*Nannospalax galili*, *Heterocephalus glaber*, *Chrysochloris asiatica*, and *Condylura cristata*) and their closest relatives in Zoonomia that do not spend time underground (*Cricetomys gambianus*, *Petromus typicus*, *Echinops telfairi*, and *Equus asinus*, respectively) using Deep SHAP version 0.39.0. We also computed the "hypothetical scores" for each nucleotide at each position for each of these sequences. We next created motifs by combining the scores and hypothetical scores using TF-MoDISco version 0.5.14.2. We visualized our per-nucleotide importance scores at OCR orthologs using normalized importance scores and the motifs using the aggregated hypothetical scores of the seqlets supporting each motif. We then created position frequency matrices by averaging the one-hot-encoded sequences at all of the seqlet coordinates belonging to the motifs. Finally, we compared each motif to known motifs by running TomTom (*125*) on the corresponding position frequency matrix with the *Homo sapiens* HOCOMOCO v11 FULL motif database (*126*).

*Evaluating the relationship between the relative rate of protein-coding gene evolution and brain size residual*

We used RERConverge to evaluate the relationship between relative evolutionary rates (RERs) of protein-coding genes and brain size residual (*19*, *127*, *128*). Specifically, we used the protein-coding gene alignments from (*129*), filtered them, and generated individual gene trees as described in (*81*). We ran RERConverge on the resulting 16,209 protein-coding gene trees using RERConverge version 0.3.0 with R version 4.1.0 (*130*), where our phenotypes were the same phenotype annotations that were used for running TACIT with brain size residual (*43*). RER-to-phenotype correlations were computed using RERconverge's correlateWithContinuousPhenotype tool, with the minimum number of species present in a gene tree for inclusion in the analysis set to 100 and the three most extreme RER and trait values in both tails winsorized (the default setting) to reduce the effects of outliers. We corrected the p-values using the Benjamini-Hochberg procedure (*45*) and considered protein-coding genes to have significant associations if their corrected p-values were less than 0.05.

*Evaluating conservation of PV+ interneuron and retina regulatory codes*

To evaluate whether the regulatory codes in PV+ interneurons and retina wer sufficiently conserved across Euarchontoglires to train CNNs that could achieve high lineage-specific OCR accuracy, we compared the TF motifs enriched for occurring in house mouse OCRs to those enriched for occurring in human OCRs. To do this, we first used bedtools v2.29 getFasta (*91*) to obtain the mm10 and hg38 sequences of the mouse and human, respectively, OCRs. We then ran MEME-ChIP (*131*) on the OCRs from each of PV+ interneurons and retina and each of house mouse and human, where we set -db to the CIS-BP version 2.00 databases (*86*) for *Mus musculus* and *Homo sapiens*, respectively, and used default settings with the exception of -meme-nmotifs 10, -spamo-skip, and -fimo-skip. After that, we compared the results from the two species.

*OCR-phenotype association pipeline*

We associated open chromatin predictions with phenotypes of interest using published R-based implementations of phylogenetic linear regression (phylolm) for continuous traits and phylogenetic logistic regression (phyloglm) for binary traits (*15*). These previously presented methods are sped-up versions of generalized least squares linear or logistic regressions in which the expected variance-covariance matrix is based on phylogenetic relationships between species (*15*, *40*, *41*). We defined the phylogenetic relationships using a phylogeny derived from the Zoonomia Cactus alignment (*132*). For both continuous and binary phenotypes, we treated the phenotype annotations as the response variable and the (continuous) probabilities of OCR ortholog open chromatin output by the CNNs as the predictor variable. For each OCR, we fit a linear or logistic model using the intersection of the species with a phenotype annotation and the species with a usable ortholog of that OCR. We conducted the association analysis for only OCRs with usable orthologs (and therefore ortholog open chromatin predictions) in at least 50% of the subset of boreoeutherian mammals with a phenotype annotation. Because, for some phenotypes, we did not have an annotation for all 222 species, the numerical threshold varied between phenotypes (**Table S11**).

We computed p-values using the published phylogenetic permulations method (*16*), in which the phenotype values are permuted in a way that produces a phenotype tree with a topology that is similar to the true phenotype tree, where the true phenotype tree was the same as the tree that we used for running phylolm and phyloglm (*132*). To reduce the computational cost, we made some modifications to the original implementation of permulations in the RERConverge package (*19*), where our modifications decreased the run-time without making any changes to the method. For permulations of both continuous and binary traits, we modified the code to allow precomputation of some phenotype-specific values that were originally computed every time a permutation was generated. We made additional more substantial speed-ups to the code for binary phenotypes because generating permulations for binary phenotypes is computationally expensive relative to doing this for continuous phenotypes, as the permulations for binary phenotypes algorithm uses a rejection sampler. From profiling, we found that most of the compute time was spent counting the number of internal nodes for which all leaf (extant species) descendents were in the foreground of a trait. This count is used for the rejection criterion. We therefore rewrote this subroutine to achieve a ~100X speedup in the counting step and a ~10X speedup overall in generating permulations of binary traits. In particular, we precompute a data structure to store which leaves are descendents of each internal node. This allows for very fast queries - using bitwise operations - of the form, "For how many internal nodes is this set of leaves [the permuted foreground] a superset of the descendants of the node?" We implemented some of this in C, using the Rcpp package (*133*).

Because of the variation in the species sets (above), we generated the permulations separately for each OCR-phenotype pair using the phenotype annotations for the specific subset of species with usable OCR orthologs. We computed the p-value for the association between an individual OCR and phenotype as the (number of trials for which the phylolm or phyloglm model fit is at least as good as the model for the real data) / (total number of trials), where we defined a trial to be a test of an association between open chromatin predictions for orthologs of an OCR and either a phenotype or a permulated phenotype. (Thus, the total number of trials is equal to the number of permulations plus one.) We considered the fit of a linear or logistic model A to be at least as good as that of another model B if the p-value that the linear term (beta) of A is non-zero is less than or equal to the p-value that the linear term (beta) of B is non-zero and if the direction of association is the same. Note that the minimum p-value possible is the reciprocal of the number of trials, as the model fit for the real data must be at least as good as itself. We corrected p-values using Benjamini-Hochberg false discovery rate (FDR) correction (*45*, *134*) with α=0.05.

Generating enough permulations to compute a precise p-value and fitting models with phylolm/phyloglm for them still had a large computational cost after implementing our speed-ups. To reduce the number of trials needed, we first performed only 1,000 trials (999 permulations) for each OCR. We then successively identified OCRs with low p-values and performed more trials (permulations) for only those OCRs. In particular, for each phenotype, we performed 9,000 more permulations (10,000 total trials) for OCRs with $p < 0.05$, then 90,000 more permulations (100,000 total trials) for OCRs with $p < 0.005$, and finally 900,000 more permulations (1,000,000 total trials) for OCRs with $p < 0.0005$. We re-computed p-values after each step. We found that the number of trials required for this staged approach was tractable, requiring only thousands to tens of thousands CPU hours for each tissue/cell type, phenotype pair. Furthermore, this approach enabled us to compute sufficiently precise p-values to evaluate statistical significance after FDR correction. In order to use p-values generated in a consistent way for comparing OCRs, we also ran 10,000 trials for all motor cortex OCRs for associations with the brain size residual phenotype and 10,000 trials for all PV+ interneuron OCRs for associations with the solitary living phenotype. We used p-values computed from these trials to compare p-values for OCRs near and not near TSSs of microcephaly and macrocephaly genes and to compare p-values for OCRs overlapping and not overlapping LD-expanded schizophrenia GWAS hits, respectively.

For running phylolm for motor cortex and liver and phyloglm for motor cortex, we used predictions for OCR orthologs in 222 boreoeutherian mammals (all boreoeutherian mammals in Zoonomia except for *Galeopterus variegatus* and *Hippopotamus amphibius*, whose chromosome names in the Cactus were not able to be mapped to chromosome names in NCBI) with usable orthologs and phenotype annotations because we had open chromatin data in both of these tissues from both Euarchontoglires and Laurasiatheria, the two clades in Boreoeutheria, but not from Afrotheria or Xenarthra, the two other mammalian clades with genomes in Zoonomia. The one exception was the associations with solitary and group living, where we used only the subset of the 222 boreoeutherian mammals that are members of the Euarchontoglires clade in order to be consistent with the analyses for PV+ interneurons, whose association with these phenotypes is supported by previous studies (*69*–*75*). For runs of our pipeline for PV+ interneurons, we used only the subset of the 222 boreoeutherian mammals that are members of the Euarchontoglires clade because we did not have any PV+ interneuron open chromatin data from Laurasiatheria. The one exception was the association with vocal learning, where we used all 222 boreoeutherian mammals because there is only one vocal learner in Euarchontoglires.

We conducted these analyses using R 4.0 (*130*). We used phylolm version 2.6 (*15*) with a Brownian motion model and default parameters as implemented in ape version 5.4.1 (*135*). We used geiger version 2.0's (*136*) sim.char method for the Brownian motion simulations underlying the permulations. We used the implementations of Benjamini-Hochberg FDR correction in IHW version 1.18 (*137*) and in the p.adjust R function (setting method = "BH").

*Connecting OCRs associated with phenotypes to relevant genes*

We identified relevant genes near OCRs associated with phenotypes by first identifying "nearby genes," which we defined as genes whose TSSs are within 1MB of the human (for OCRs initially identified in rhesus macaque or human) or house mouse (for OCRs initially identified in house mouse or Norway rat) orthologs of these OCRs. For OCRs initially identified in Egyptian fruit bat, we used human TSSs for OCRs with a usable human ortholog and mouse TSSs for other OCRs. We defined human TSSs as the protein-coding TSSs annotated in human GENCODE version 27 and mouse TSSs as the protein-coding TSSs annotated in house mouse GENCODE version 15 (*106*, *138*). We used OMIM (*47*) to determine which of the nearby genes have protein-coding mutations associated with disorders related to the associated phenotype. To identify the full set of motor cortex OCRs near microcephaly and macrocephaly genes, we first identified a list of disorders whose OMIM clinical synopses included one of the following search terms: "microcephaly," "macrocephaly," "small head circumference," or "large head circumference," excluding entries describing common variation. We next restricted this list of disorders to those caused by one or more known, single-gene mutations. We then obtained the regions within 1Mb of these genes' TSSs and identified all overlapping motor cortex human OCR orthologs that were tested for association with brain size residual. We compared the p-values of this set of OCRs to the p-values of all of the remaining motor cortex OCRs with usable human orthologs that were tested for associations with brain size residual. We used a one-sided Wilcoxon rank-sum test, where our null hypothesis was that there was no difference in the distributions of p-values and our alternative hypothesis was that the p-values for OCRs with nearby microcephaly and macrocephaly genes were lower than the p-values for the other OCRs.

*Evaluating 3D connections between human motor cortex OCR orthologs and genes related to brain size*

We investigated whether human orthologs of motor cortex OCRs whose open chromatin is positively associated with brain size residual are close in 3D space to nearby relevant genes using adult cortex Hi-C data from (*55*). To match the assembly used in the Hi-C data, we used liftOver (*108*) to map these orthologs (**Table S8**) from hg38 to hg19. We then used HUGIn (http://hugin2.genetics.unc.edu/Project/hugin/) to obtain 3D genome interactions from this dataset. Specifically, we considered each gene as an anchor and obtained the p-value for the association between the bin containing each OCR ortholog and that gene. We excluded OCRs without a human ortholog, OCRs on the X chromosome, and genes not found in the anchor database of HUGIn. In total, we investigated 24 gene-OCR pairs that included 12 OCRs (**Table S8**). We corrected those p-values using Benjamini-Hochberg FDR correction (*45*, *134*).

*Comparing associations of PV+ interneuron OCRs with solitary living for PV+ interneuron OCRs with human orthologs overlapping GWAS hits to other PV+ interneuron OCRs*

We obtained genetic variants associated with schizophrenia from a schizophrenia GWAS (*77*) and used all variants with $p < 5 \times 10^{-8}$. We next obtained all human single-nucleotide polymorphisms with linkage disequilibrium > 0.5 with the GWAS variants (*139*) and combined these sets of variants. We then mapped the combined variant set from hg19 to hg38 using liftOver (*108*). After that, we identified all human orthologs of PV+ interneuron OCRs used for solitary living associations that did and did not overlap the combined set of variants using bedtools (*91*). Finally, we compared the p-value distributions for each group of OCRs, using p-values generated by applying Cell-TACIT genome-wide using 10,000 permulations for each OCR, with a 1-sided Wilcoxon rank-sum test, where our null hypothesis was that there was no difference in the distributions of p-values and our alternative hypothesis was that the p-values for OCRs overlapping variants from our combined set were lower than the p-values for the other OCRs with human orthologs. We did not repeat this analysis for group living because our non-group living phenotype included both solitary living and monogamous species, and we expected that both group living and monogamy might be negatively associated with schizophrenia.

#### Supplementary Text

*Interpreting bulk tissue CNNs yielded relevant TF motifs*

We interpreted each CNN trained on motor cortex and liver data to evaluate if the CNNs had learned motifs of TFs that are known to be involved in the tissue in which they were trained. When interpreting the mouse-only motor cortex CNN, we found that it seemed to have learned the motifs for multiple TFs known to be involved in the motor cortex, including the motifs of Egr3 (*140*), Neurod2 (*141*), Mef2c (*32*,

*33*), Jund (*142*), Ctcf (*143*, *144*), and Rfx4 (*145*, *146*), as well as the motifs of Ascl2 and Zfp287 and some motifs that did not match any known TF motifs (**Fig. S5**). When interpreting the multi-species motor cortex CNN, we found motifs of many of the TFs with known roles in the motor cortex that we found when interpreting the mouse-only motor cortex CNN, including the motifs of EGR3, MEF2C, NEUROD2, CTCF, and RFX4, as well as additional motifs of TFs known to be involved in the motor cortex, including the motifs of NFE2L1 (*147*), HLF (*148*), SOX4 (*149*, *150*), TBR1 (*151*), TBP (*152*), and LHX2 (*153*, *154*), and some motifs that did not match any known TF motifs. For the multi-species motor cortex CNN, we also found a depletion of the motifs for ONECUT1, NR2F6, and ZNF235, all of which have low expression in the human cortex (*59*, *155*), and of the motif for ZNF785, which might be explained by a depletion of repetitive elements (**Fig. S6**). When interpreting the multi-species liver CNN, we found the motifs of many TFs known to be involved in the liver, including RXRA (*156*, *157*), FOXA3 (*158*, *159*), HNF4A (*34*, *35*), CEBPA (*160*, *161*), and ZNF529 (*162*), as well as some motifs that did not match any known TF motifs (**Fig. S7**). These results suggest that our CNNs are learning sequence patterns that are relevant to the tissue for which we trained them to predict open chromatin.

*Conservation of the PV+ interneuron regulatory code across Euarchontoglires*

To evaluate the conservation of the PV+ interneuron regulatory code between house mouse and human, we identified motifs enriched for occurring in the OCRs for each species and compared our findings. We found an enrichment of motifs for multiple TFs in peaks from both species, including the motifs for Ctcf, which has been shown to be involved in anchoring DNA loops across cell types and metazoans (*163*); Mef2d, which has been shown to play a role in neuronal activity-dependent gene regulation (*60*, *164*); Atf3, which has been shown to be activated in injured PV+ interneurons (*165*); Etv1, which has been shown to help specify the PV+ interneuron lineage (*166*); Maf, which has been shown to promote PV+ interneuron cell fate in the medial ganglionic eminence (*167*); Atf7; Junb; Nhlh2; Zbtb6; Zbtb37; Myrf;

Plagl2; Tcfl5; and Zbtb3 (**Supplementary Website**). The fact that motifs of multiple TFs, including TFs known to be involved in PV+ interneurons, are enriched for occurring in OCRs from both house mouse and human suggest that the regulatory code of PV+ interneurons is at least partially conserved between Glires and Euarchonta.

*Partial conservation of the retinal regulatory code across Euarchontoglires*

We used the same process that we used for PV+ interneurons to evaluate the conservation of the retina regulatory code between house mouse and human. We found an enrichment of motifs for multiple TFs in peaks from both species, including the motifs for Ctcf, which has been shown to be involved in anchoring DNA loops across cell types and metazoans (*163*); Vax2, which has been shown to play a role in retinal patterning (*168*) and be highly expressed in the adult primate retina (*169*); Smad4, which has been shown to be involved in angiogenesis in the retina (*170*); Hic1, whose expression is associated with age-related macular degeneration (*171*); Zfx, whose expression in the retina increases in response to optic nerve injury (*172*); Cdc5l; Ascl2; Zic1; Zbtb3; Zbtb6; Nfic; Bcl11a; Foxn1; Znf449; Znf770; Zbtb37; Rest; Znf692; Plagl1; Mafb; Thap12; Zbtb7a; Msantd3; Insm1; Ovol2; Zbtb49; Znf771; Nfkb2; Znf382; Znf189 and Cenpb. However, the motifs for CRX and NR2E3, which have been shown to play important roles in retinal cell fate determination (*173*), were enriched for occurring in only the human OCRs, suggesting that either the retinal regulatory code is not fully conserved between mouse and human or that the mouse and human data had substantial differences in retinal cell type representation (**Supplementary Website**).

*Species cluster by clade when using predicted open chromatin as features*

We investigated whether there is a relationship between species' motor cortex OCR orthologs' open chromatin predictions and species' locations on the phylogenetic tree. **Fig. S4** shows a boreoeutherian mammal tree constructed by hierarchical clustering of the open chromatin predictions of all motor cortex OCR orthologs in each species. In general, species cluster by clade. Of the 20 clades shown, only two small clades (Euliopotyphyla and Casotorimorpha) do not appear as monophyletic in the tree. At a high level, the reconstruction of the high-level Laurasiathera tree is very similar to the established phylogeny, as is the Primate subtree (*132*). Interestingly, the clades that cluster in incorrect phylogenetic positions – outside of all other mammals in the tree – are the muroid rodents and the shrews, which have exceptionally fast inter-generation times compared to all other mammals that could explain their faster rates of genome evolution compared to other mammals (*174*, *175*). These faster genome evolutionary rates may also be associated with faster changes in motor cortex open chromatin.  In addition, we have data from two muroid rodents and zero or one member of each other clade, so muroid rodents may generally have higher predicted activity than other clades.

*TACIT identifies liver OCRs associated with the evolution of brain size*

We also identified liver OCRs whose predicted open chromatin status is associated with brain size residual. We found 73 such OCRs after FDR correction, of which 9 were negatively associated and 64 were positively associated (**Table S12**). None of these overlap motor cortex or PV+ interneuron OCRs that are significantly associated with brain size residual. We may have found different liver OCRs associated with brain size residual because the liver is involved in non-neurological processes associated with brain size, such as metabolism.

**Supplementary Figures**

**Figure S1: Mouse-only CNN evaluations**

**(A-C)** show the AUROC and the AUPRC/NPV-Spec. for the full test set, for OCRs and non-OCR orthologs of OCRs in non-house mouse species, and for lineage-specific OCR accuracy evaluations for mouse-only motor cortex (**A**), PV+ interneuron (**B**), and retina (**C**) CNNs. **(D-F)** show the AUROC and AUPRC/NPV-Spec. for tissue-specific OCR accuracy evaluations for mouse-only motor cortex (**D**), PV+ interneuron (**E**), and retina (**F**) CNNs. **(G-L)** show the negative relationship between the average house mouse OCR ortholog mouse-only motor cortex (**G** and **J**), PV+ interneuron (**H** and **K**), and retina (**I** and **L)** CNN predictions for Glires species and the MYA when species diverged from house mouse, where (**G-I**) are from after removing OCR orthologs for which >5% of the nucleotides are N's and (**J-L**) are from everything. **(M-R)** show the positive relationship between the standard deviation of house mouse OCR ortholog mouse-only motor cortex (**M** and **P**), PV+ interneuron (**N** and **Q**), and retina (**O** and **R**) CNN predictions for Glires species and the MYA when species diverged from house mouse, where (**M-O**) are from after removing OCR orthologs for which >5% of the nucleotides are N's, and (**P-R**) are from everything.

**Figure S2: Additional motor cortex, PV+ interneuron, and retina multi-species CNN phylogeny-matching correlations evaluations**

**(A-B)** show the negative relationship between the average house mouse OCR ortholog multi-species motor cortex (**A**) and PV+ interneuron (**B**) CNN predictions for Glires species and the MYA when species diverged from house mouse, using predictions from everything (removing sequences with >5% Ns is displayed in **Fig. 2**). **(C-F)** show the positive relationship between the standard deviation of house mouse OCR ortholog multi-species motor cortex (**C** and **E**) and PV+ interneuron (**D** and **F**) CNN predictions for Glires species and the MYA when species diverged from house mouse, where (**C-D**) are from removing OCR orthologs for which >5% of the nucleotides are N's and (**E-F**) are from everything.

**Figure S3: Liver multi-species CNN evaluations**

**(A)** shows the AUROC and the area under the AUPRC/NPV-Spec. for the full test set, clade-specific OCRs and non-OCRs, and shared versus tissue-specific OCRs and non-OCRs for multi-species liver CNNs. **(B-C)** show the negative relationship between the average house mouse OCR ortholog multi-species liver CNN predictions for Glires species and the MYA when species diverged from house mouse, where (**B**) is from removing OCR orthologs for which more 5% of the nucleotides are N's and (**C**) is from everything. **(D-E)** show the positive relationship between the standard deviation of house mouse OCR ortholog multi-species liver CNN predictions for Glires species and the MYA when species diverged from house mouse, where (**D**) is from removing OCR orthologs for which more 5% of the nucleotides are N's, and (**E**) is from everything.

**Figure S4: Dendrogram of boreoeutherian mammal relationships inferred from motor cortex OCR ortholog open chromatin predictions**

A dendrogram of species inferred by average-linkage clustering of species, with pairwise distances computed by taking 1 minus the cosine similarity between the vectors of motor cortex OCR ortholog open chromatin predictions, using for each pair of species the OCRs with orthologs in both species. Clustering was done using the hclust function in R. Species are grouped into 20 clades and colored accordingly; clade names are shown at the right. Closely related clades are generally similar colors, e.g. all the Euarchonta clades are different shades of green. Two key points in the tree that are inferred correctly, the roots of Laurasiatheria and Primates, are indicated.

**Figure S5: TF-MoDISco results for mouse-only motor cortex CNN**

Motifs from running TF-MoDISco on the positive validation set with the mouse-only motor cortex CNN, transcription factors (TFs) whose motifs match the TF-MoDISco motifs with TomTom q-value < 0.05 ordered from most to least significant TomTom p-value (*125*), and number of supporting seqlets for each motif. Underlined TFs are those whose roles in the motor cortex are highlighted in the **Supplementary Text**.

**Figure S6: TF-MoDISco results for multi-species motor cortex CNN**

Motifs from running TF-MoDISco on the positive validation set with the multi-species motor cortex CNN; transcription factors (TFs) whose motifs match the TF-MoDISco motifs with TomTom q-value < 0.05 ordered from most to least significant TomTom p-value (*125*), where red TFs are those whose motifs are considered important and gold TFs are those whose motifs' depletions are considered important; and number of supporting seqlets for each motif. Underlined TFs are those whose roles in the motor cortex are highlighted in the **Supplementary Text**.

**Figure S7: TF-MoDISco results for multi-species liver CNN**

Motifs from running TF-MoDISco on the positive validation set with the multi-species liver CNN, transcription factors (TFs) whose motifs match the TF-MoDISco motifs with TomTom q-value < 0.05 ordered from most to least significant TomTom p-value (*125*), and number of supporting seqlets for each motif. Underlined TFs are those whose roles in the liver are highlighted in the **Supplementary Text**.

**Figure S8: Additional examples of motor cortex and PV+ interneuron OCRs associated with brain size**

Plots of brain size residual versus predicted OCR ortholog open chromatin probability for eight motor cortex **(A-H)** OCRs associated with brain size residual in boreoeutherian mammals and three PV+ interneuron **(I-K)** OCRs associated with brain size residual in Euarchontoglires. Each OCR is within 1Mb of a gene mutated in macro- or microcephaly as indicated at the top of each panel; see **Tables S8-S9** for lists of all brain size-relevant genes near each OCR. Points are colored by clade, and shapes highlight key lineages as indicated. Lines show the phylogeny-corrected lines of best fit from phylolm, both overall (solid black) and within each clade (dashed). Coordinates reported are those of the OCR in the species in which it was originally identified. For the two pairs of OCRs both near *Satb1*/*SATB1* and *Mocs2*, the distinguishing numbers (1) and (2) match those used in **Figs. 4** and **5**, respectively.

**Figure S9: Boreoeutherian mammal-wide associations between predicted motor cortex OCR ortholog open chromatin and brain size residual of OCRs in the SATB1 locus**

**(A-B)** show the Boreoeutheria-wide versions of **Figs. 3A-B**: predicted motor cortex open chromatin versus brain size residual of orthologs of two motor cortex OCRs in the SATB1 locus, chr17:52351209-52351928 (mm10) and chr2:174466184-174466517 (rheMac8). Each point represents one ortholog, and points are grouped along the x-axis of each panel by clade as shown by the tree below. The clades and example species are listed in **Table S10**. Points are colored by brain size residual following the scale at the bottom. The permulations p-value after Benjamini-Hotchberg correction and the coefficient on the predicted open chromatin returned by phylolm are shown in the lower right of each panel.

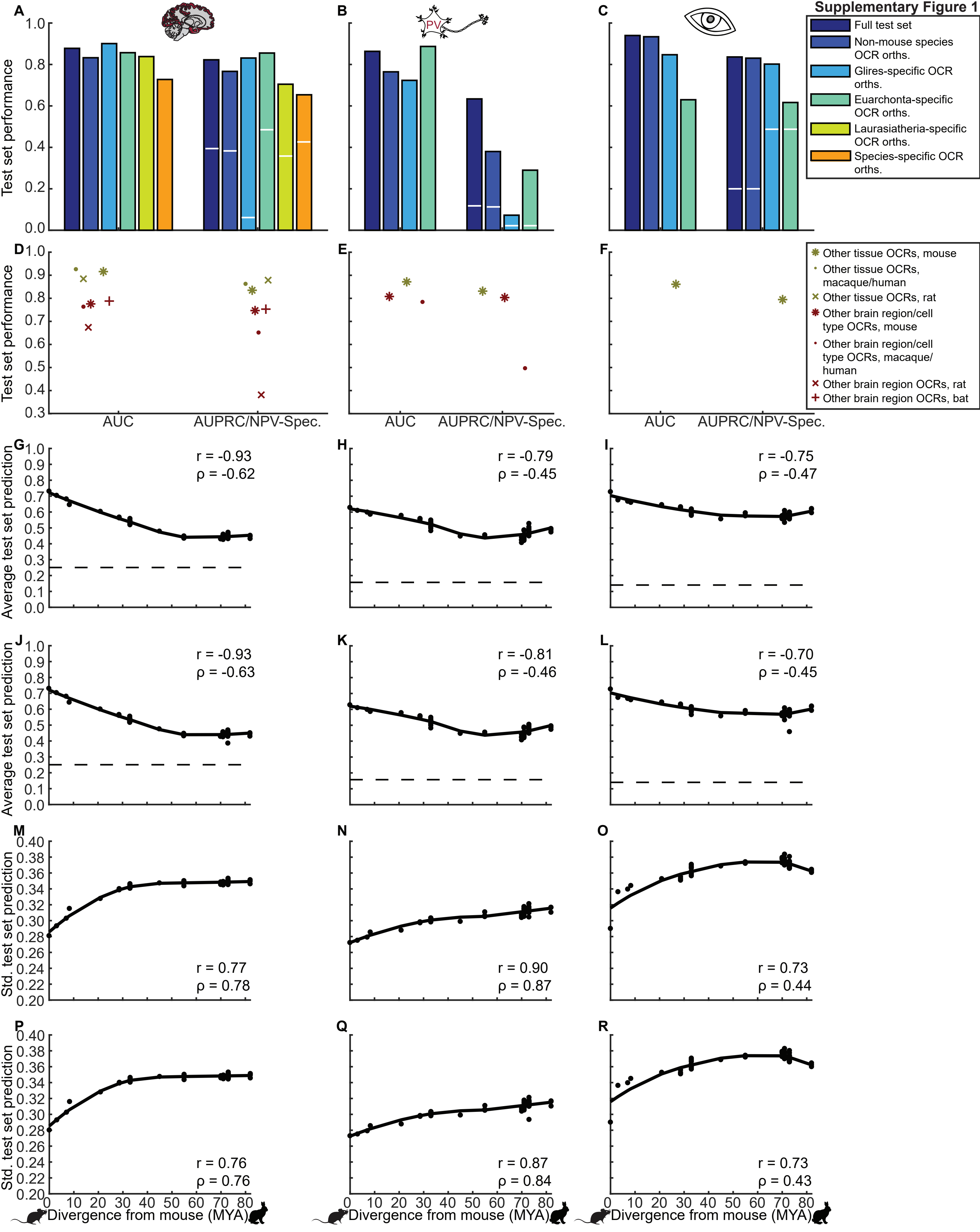

Supplementary Figure 1
A
B
C
PV
Full test set
Non-mouse species OCR orths.
Glires-specific OCR orths.
Euarchonta-specific OCR orths.
Laurasiatheria-specific OCR orths.
Species-specific OCR orths.
Test set performance
D
E
F
Other tissue OCRs, mouse
Other tissue OCRs, macaque/human
Other tissue OCRs, rat
Other brain region/cell type OCRs, mouse
Other brain region/cell type OCRs, macaque/ human
Other brain region OCRs, rat
Other brain region OCRs, bat
AUC
AUPRC/NPV-Spec.
G
H
I
r = -0.93
ρ = -0.62
r = -0.79
ρ = -0.45
r = -0.75
ρ = -0.47
Average test set prediction
J
K
L
r = -0.93
ρ = -0.63
r = -0.81
ρ = -0.46
r = -0.70
ρ = -0.45
M
N
O
r = 0.77
ρ = 0.78
r = 0.90
ρ = 0.87
r = 0.73
ρ = 0.44
Std. test set prediction
P
Q
R
r = 0.76
ρ = 0.76
r = 0.87
ρ = 0.84
r = 0.73
ρ = 0.43
Divergence from mouse (MYA)

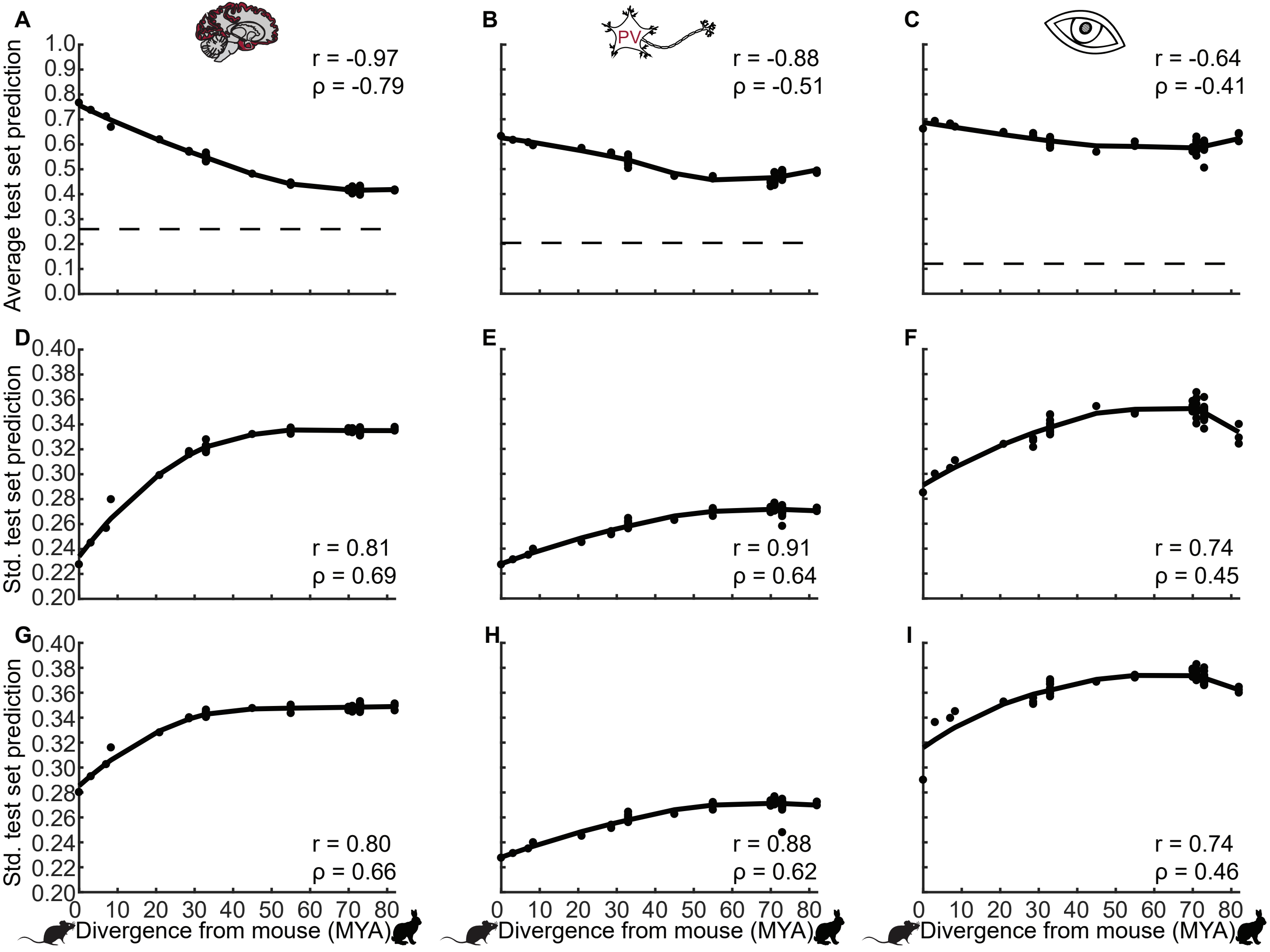
A
B
C
PV
r = -0.97
ρ = -0.79
r = -0.88
ρ = -0.51
r = -0.64
ρ = -0.41
Average test set prediction
1.0
0.9
0.8
0.7
0.6
0.5
0.4
0.3
0.2
0.1
0.0
D
E
F
r = 0.81
ρ = 0.69
r = 0.91
ρ = 0.64
r = 0.74
ρ = 0.45
Std. test set prediction
0.40
0.38
0.36
0.34
0.32
0.30
0.28
0.26
0.24
0.22
0.20
G
H
I
r = 0.80
ρ = 0.66
r = 0.88
ρ = 0.62
r = 0.74
ρ = 0.46
Std. test set prediction
0 10 20 30 40 50 60 70 80
Divergence from mouse (MYA)
Divergence from mouse (MYA)
Divergence from mouse (MYA)

#### Supplementary Figure 3

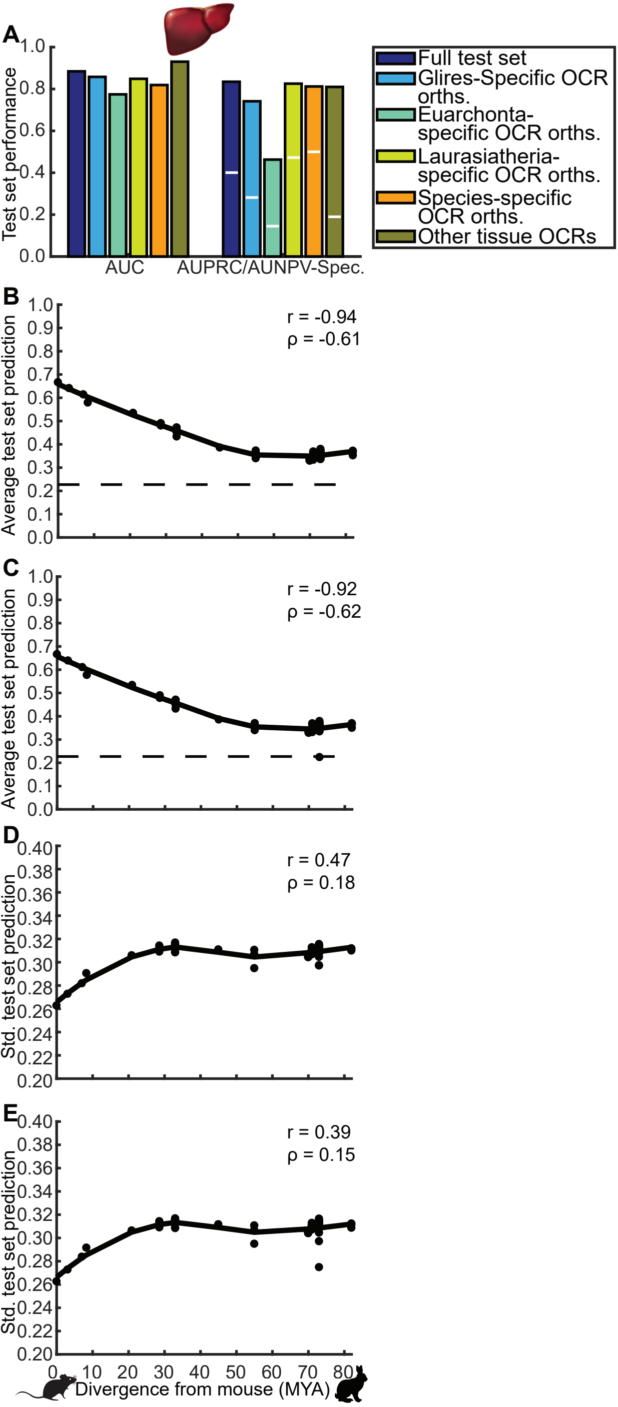

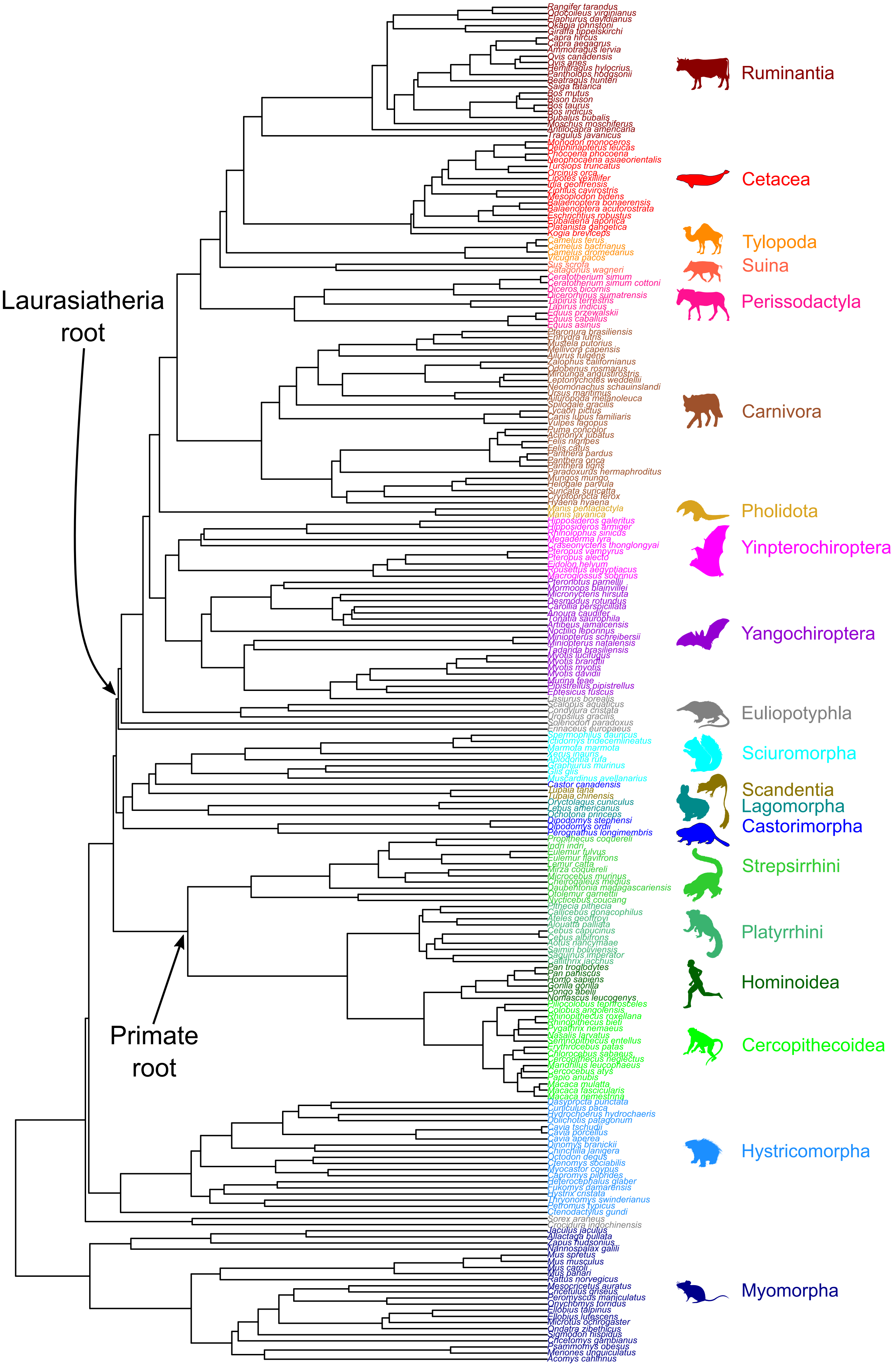
Laurasiatheria root
Primate root
Rangifer tarandus
Odocoileus virginianus
Elaphurus davidianus
Okapia johnstoni
Giraffa tippelskirchi
Capra hircus
Capra aegagrus
Ammotragus lervia
Ovis canadensis
Ovis aries
Hemitragus hylocrius
Pantholops hodgsonii
Beatragus hunteri
Saiga tatarica
Bos mutus
Bison bison
Bos taurus
Bos indicus
Bubalus bubalis
Moschus moschiferus
Antilocapra americana
Tragulus javanicus
Monodon monoceros
Delphinapterus leucas
Phocoena phocoena
Neophocaena asiaeorientalis
Tursiops truncatus
Orcinus orca
Lipotes vexillifer
Inia geoffrensis
Ziphius cavirostris
Mesoplodon bidens
Balaenoptera bonaerensis
Balaenoptera acutorostrata
Eschrichtius robustus
Eubalaena japonica
Platanista gangetica
Kogia breviceps
Camelus ferus
Camelus bactrianus
Camelus dromedarius
Vicugna pacos
Sus scrofa
Catagonus wagneri
Ceratotherium simum
Ceratotherium simum cottoni
Diceros bicornis
Dicerorhinus sumatrensis
Tapirus terrestris
Tapirus indicus
Equus przewalskii
Equus caballus
Equus asinus
Pteronura brasiliensis
Enhydra lutris
Mustela putorius
Mellivora capensis
Ailurus fulgens
Zalophus californianus
Odobenus rosmarus
Mirounga angustirostris
Leptonychotes weddellii
Neomonachus schauinslandi
Ursus maritimus
Ailuropoda melanoleuca
Spilogale gracilis
Lycaon pictus
Canis lupus familiaris
Vulpes lagopus
Puma concolor
Acinonyx jubatus
Felis nigripes
Felis catus
Panthera pardus
Panthera onca
Panthera tigris
Paradoxurus hermaphroditus
Mungos mungo
Helogale parvula
Suricata suricatta
Cryptoprocta ferox
Hyaena hyaena
Manis pentadactyla
Manis javanica
Hipposideros galeritus
Hipposideros armiger
Rhinolophus sinicus
Megaderma lyra
Craseonycteris thonglongyai
Pteropus vampyrus
Pteropus alecto
Eidolon helvum
Rousettus aegyptiacus
Macroglossus sobrinus
Pteronotus parnellii
Mormoops blainvillei
Micronycteris hirsuta
Desmodus rotundus
Carollia perspicillata
Anoura caudifer
Tonatia saurophila
Artibeus jamaicensis
Noctilio leporinus
Miniopterus schreibersii
Miniopterus natalensis
Tadarida brasiliensis
Myotis lucifugus
Myotis brandtii
Myotis myotis
Myotis davidii
Murina feae
Pipistrellus pipistrellus
Eptesicus fuscus
Lasiurus borealis
Scalopus aquaticus
Condylura cristata
Uropsilus gracilis
Solenodon paradoxus
Erinaceus europaeus
Spermophilus dauricus
Ictidomys tridecemlineatus
Marmota marmota
Xerus inauris
Aplodontia rufa
Graphiurus murinus
Glis glis
Muscardinus avellanarius
Castor canadensis
Tupaia tana
Tupaia chinensis
Oryctolagus cuniculus
Lepus americanus
Ochotona princeps
Dipodomys stephensi
Dipodomys ordii
Perognathus longimembris
Propithecus coquereli
Indri indri
Eulemur fulvus
Eulemur flavifrons
Lemur catta
Mirza coquereli
Microcebus murinus
Cheirogaleus medius
Daubentonia madagascariensis
Otolemur garnettii
Nycticebus coucang
Pithecia pithecia
Callicebus donacophilus
Ateles geoffroyi
Alouatta palliata
Cebus capucinus
Cebus albifrons
Aotus nancymaae
Saimiri boliviensis
Saguinus imperator
Callithrix jacchus
Pan troglodytes
Pan paniscus
Homo sapiens
Gorilla gorilla
Pongo abelii
Nomascus leucogenys
Piliocolobus tephrosceles
Colobus angolensis
Rhinopithecus roxellana
Rhinopithecus bieti
Pygathrix nemaeus
Nasalis larvatus
Semnopithecus entellus
Erythrocebus patas
Chlorocebus sabaeus
Cercopithecus neglectus
Mandrillus leucophaeus
Cercocebus atys
Papio anubis
Macaca mulatta
Macaca fascicularis
Macaca nemestrina
Dasyprocta punctata
Cuniculus paca
Hydrochoerus hydrochaeris
Dolichotis patagonum
Cavia tschudii
Cavia porcellus
Cavia aperea
Dinomys branickii
Chinchilla lanigera
Octodon degus
Ctenomys sociabilis
Myocastor coypus
Capromys pilorides
Heterocephalus glaber
Fukomys damarensis
Hystrix cristata
Thryonomys swinderianus
Petromus typicus
Ctenodactylus gundi
Sorex araneus
Crocidura indochinensis
Jaculus jaculus
Allactaga bullata
Zapus hudsonius
Nannospalax galili
Mus spretus
Mus musculus
Mus caroli
Mus pahari
Rattus norvegicus
Mesocricetus auratus
Cricetulus griseus
Peromyscus maniculatus
Onychomys torridus
Ellobius lutescens
Microtus ochrogaster
Ondatra zibethicus
Sigmodon hispidus
Cricetomys gambianus
Psammomys obesus
Meriones unguiculatus
Acomys cahirinus
Ruminantia
Cetacea
Tylopoda
Suina
Perissodactyla
Carnivora
Pholidota
Yinpterochiroptera
Yangochiroptera
Eulipotyphla
Sciuromorpha
Scandentia
Lagomorpha
Castorimorpha
Strepsirrhini
Platyrrhini
Hominoidea
Cercopithecoidea
Hystricomorpha
Myomorpha

| Cortex TF-MoDISco motifs | TFs with similar motifs | Seqlets |
|---|---|---|
| 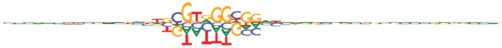 | Rela, Bcl6, Egr3 | 876 |
| 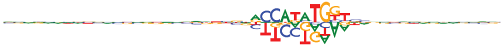 | Bhlhe22, Atoh1, Neurod2, Neurog1, Msc, Twist2, Tcf21, Atoh7, Olig1, Olig2, Bhlhe23, Bhlha15, Ascl2, Ferd3l, Neurog2, Tfap4, Twist1 | 626 |
| 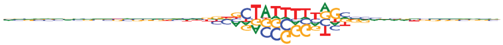 | Mef2a, Mef2c, Zfp287 | 617 |
| 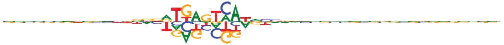 | Jund | 550 |
| 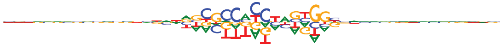 | Ctcf, Zfp661 | 435 |
| 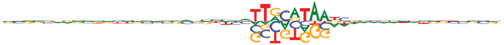 | | 180 |
| 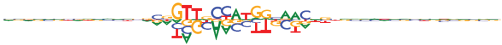 | Rfx1, Rfx4, Rfx5, Rfx2, Rfx7, Rfx3 | 174 |
| 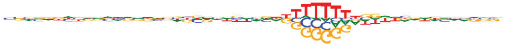 | Zfp287, Zfp422, Prdm6, Zfp384 | 149 |
| 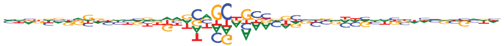 | Ascl2 | 62 |
| 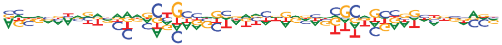 | | 26 |

#### Supplementary Figure 6

| Cortex TF-MoDISco motifs | TFs with similar motifs | Seqlets |
|---|---|---|
| 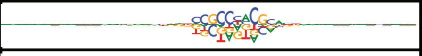 | ZNF432, EGR3, EGR1, ZNF212, ZNF467, ZSCAN22, EGR4 | 2192 |
| 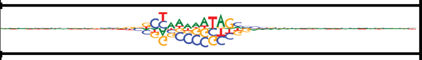 | MEF2A, ZNF235, MEF2D, ZNF287, MEF2B, MEF2C | 1974 |
| 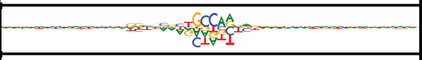 | | 1261 |
| 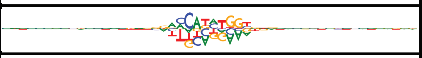 | OLIG3, ZNF287, BHLHE23, OLIG2, ZNF235, OLIG1, BHLHA15, NEUROD2, ATOH7, BHLHE22, ZNF148, NEUROG1, NEUROG2, TWIST1, ZFP28, NEUROD1, FIGLA | 1205 |
| 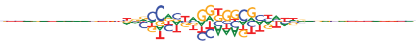 | CTCF, CTCFL | 1103 |
| 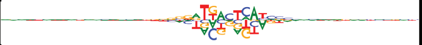 | NFE2L1, JUN, JDP2, MAFG, BACH2, NFE2, MAFB | 770 |
| 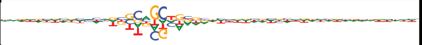 | | 674 |
| 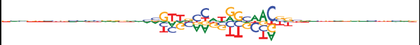 | RFX3, RFX6, RFX5, RFX1, RFX4, NFATC1, RFX8 | 354 |
| 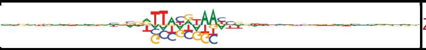 | ZNF235, HLF, NFIL3, TEF, CEBPE, MITF, CENPBD1, ZNF287, DBP, FOXQ1 | 280 |
| 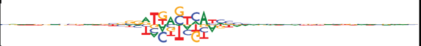 | | 261 |
| 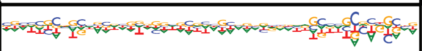 | | 175 |
| 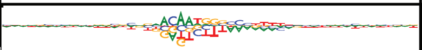 | SOX4, SOX9 | 119 |
| 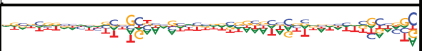 | | 109 |
| 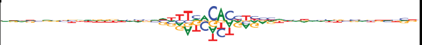 | TBX18, TBX22, TBR1, TBX10, TBX6, EOMES, TBX3 | 107 |
| 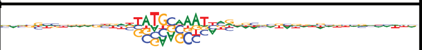 | POU2F3, POU3F1, POU3F3, TBP | 79 |
| 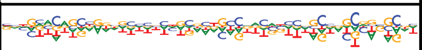 | | 66 |
| 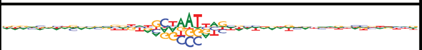 | ALX3, VSX1, ESX1, LHX2, NKX1-2, HESX1, PROP1, LHX8, OTP, SHOX2, EMX2, RAX2, NOTO, ARGFX, LHX9, HOXD8, EVX2, MIXL1, RAX, ZFHX3, EVX1, MSX2, PRRX2, SHOX, GBX1, PRRX1, EN1, HOXA1, HOXD3, HOXD1, POU3F1, POU3F3, ZNF655, VAX2, SEBOX, HOXB3, BSX, GBX2, HOXB2, ISX, NKX1-1, EMX1, LBX1, GSX2, MNX1, GSX1, ARX, HOXA5, LHX5, LHX4, EN2, LHX3 | 47 |
| 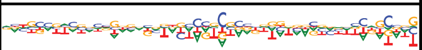 | | 34 |
| 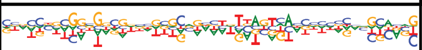 | | 25 |
| 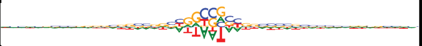 | ZNF235, ZNF287, ONECUT3, ONECUT1, ONECUT2, HMG20B, DBX1, TLX1, PBX4, LHX3 | 876 |
| 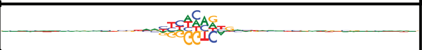 | NR2F6 | 855 |
| 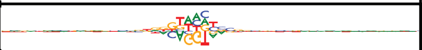 | ZNF235, ZNF287, ZEB1 | 756 |
| 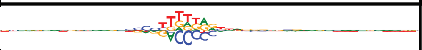 | | 185 |
| 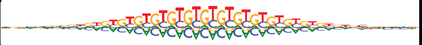 | ZNF785 | 32 |

**Supplementary Figure 7**

| Liver TF-MoDISco motifs | TFs with similar motifs | Seqlets |
|---|---|---|
| 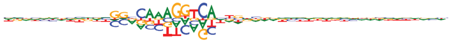 | RXRA, RXRB, NR2C2, HNF4G, RXRG, PPARD, NR2F1, NR4A3, NR2E1, RARB | 252 |
| 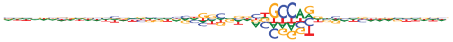 | | 237 |
| 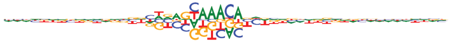 | ZNF287, FOXR1, FOXB2, ZNF235, FOXJ1, FOXP3, PRDM6, FOXF1, FOXA3, FOXE3, FOXE1, FOXL1, FOXS1, FOXN3, FEZF2, FOXI1, FOXQ1, FOXI2 | 196 |
| 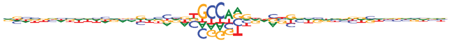 | | 154 |
| 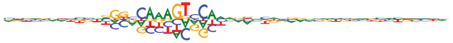 | HNF4G, HNF4A, NR2C2, NR2F1, PPARD, RXRA, RARB | 103 |
| 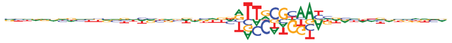 | CEBPB, CEBPE, CEBPD, CEBPG, CEBPA, ZBTB22, HLF, DBP, TEF | 92 |
| 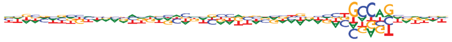 | | 74 |
| 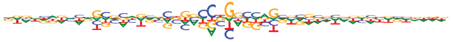 | | 83 |
| 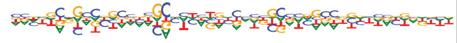 | | 62 |
| 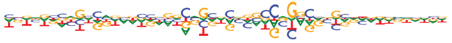 | | 62 |
| 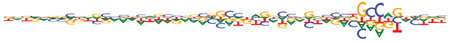 | | 50 |
| 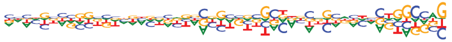 | | 41 |
| 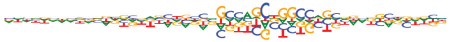 | | 57 |
| 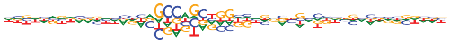 | | 41 |
| 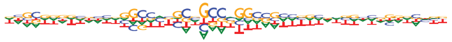 | ZNF180, ZNF792, ZIM2, ZNF529, ZNF283, ZNF304, ZNF341 | 56 |
| 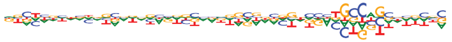 | | 28 |
| 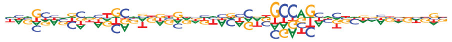 | | 34 |
| 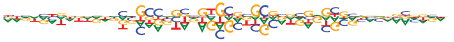 | | 28 |

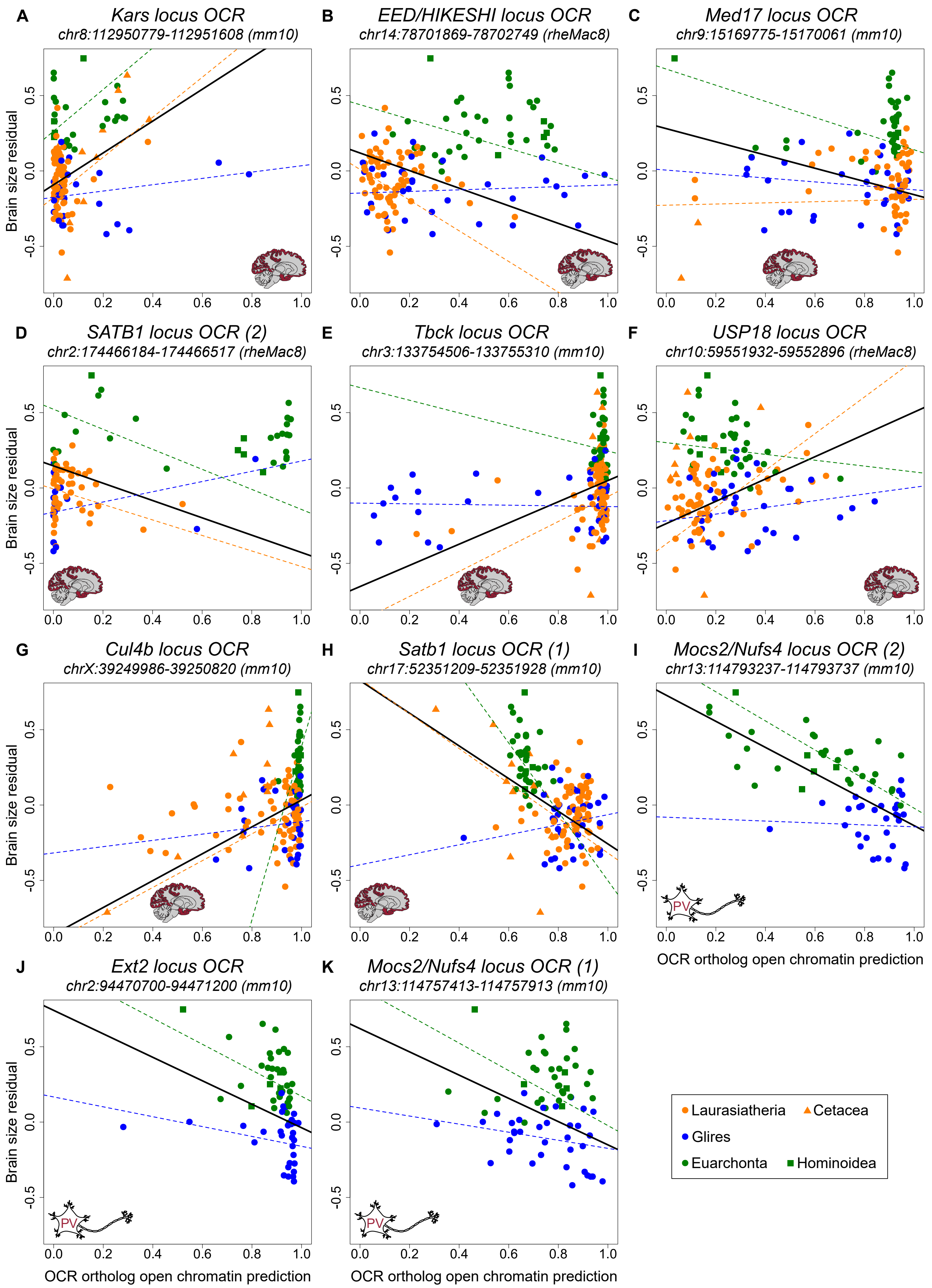
A
Kars locus OCR
chr8:112950779-112951608 (mm10)
B
EED/HIKESHI locus OCR
chr14:78701869-78702749 (rheMac8)
C
Med17 locus OCR
chr9:15169775-15170061 (mm10)
D
SATB1 locus OCR (2)
chr2:174466184-174466517 (rheMac8)
E
Tbck locus OCR
chr3:133754506-133755310 (mm10)
F
USP18 locus OCR
chr10:59551932-59552896 (rheMac8)
G
Cul4b locus OCR
chrX:39249986-39250820 (mm10)
H
Satb1 locus OCR (1)
chr17:52351209-52351928 (mm10)
I
Mocs2/Nufs4 locus OCR (2)
chr13:114793237-114793737 (mm10)
J
Ext2 locus OCR
chr2:94470700-94471200 (mm10)
K
Mocs2/Nufs4 locus OCR (1)
chr13:114757413-114757913 (mm10)
Brain size residual
OCR ortholog open chromatin prediction
PV
Laurasiatheria
Cetacea
Glires
Euarchonta
Hominoidea

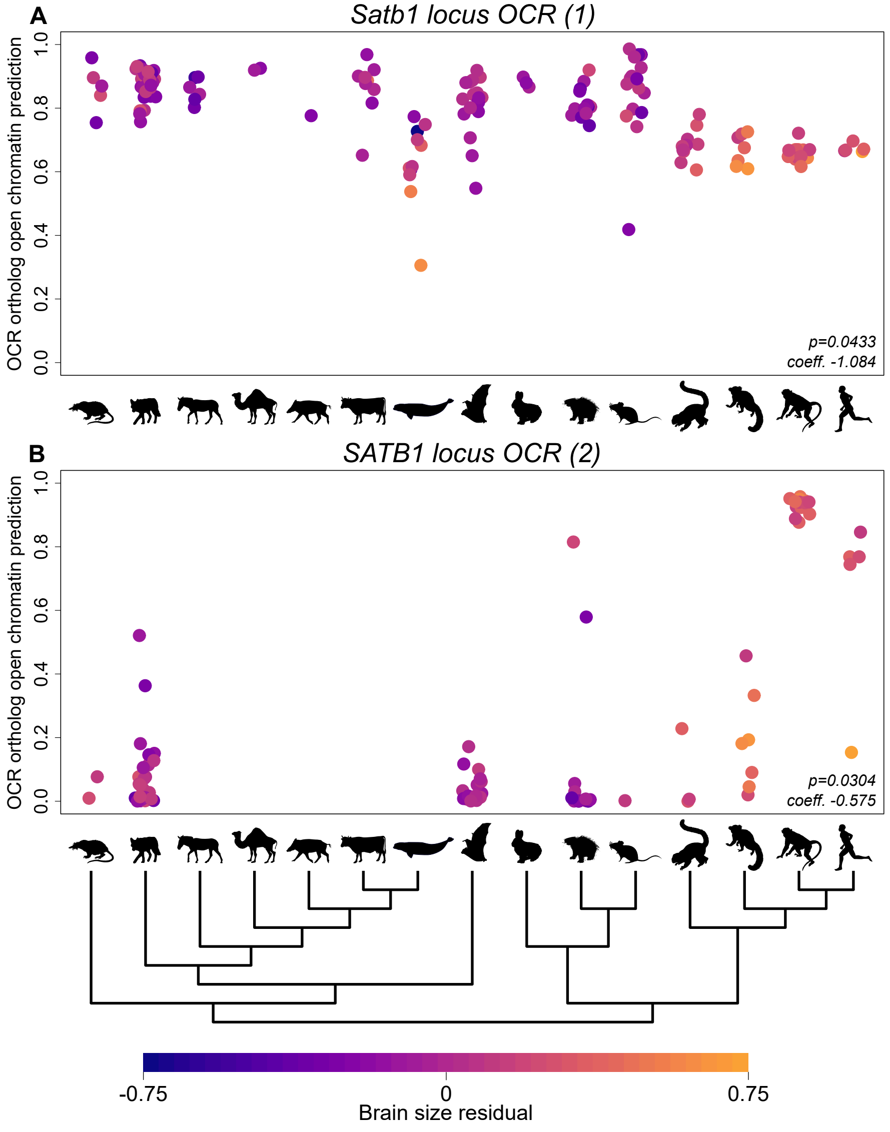
A
Satb1 locus OCR (1)
OCR ortholog open chromatin prediction
0.0
0.2
0.4
0.6
0.8
1.0
p=0.0433
coeff. -1.084
B
SATB1 locus OCR (2)
OCR ortholog open chromatin prediction
0.0
0.2
0.4
0.6
0.8
1.0
p=0.0304
coeff. -0.575
-0.75
0
0.75
Brain size residual

**Supplementary Tables**

| Evaluation Subset of Test Chromosome Orthologs | Fraction of Minority Class | Sensitivity | Specificity | Precision | NPV |
|---|---|---|---|---|---|
| Full Test Set | 0.39 | 0.78 | 0.82 | 0.87 | 0.70 |
| Full Test Set from Non-Mouse Species | 0.38 | 0.75 | 0.76 | 0.84 | 0.66 |
| Glires-Specific OCRs and Non-OCRs | 0.061 | 0.79 | 0.85 | 0.91 | 0.68 |
| Euarchonta-Specific OCRs and Non-OCRs | 0.49 | 0.78 | 0.78 | 0.79 | 0.77 |
| Laurasiatheria-Specific OCRs and Non-OCRs | 0.36 | 0.83 | 0.72 | 0.62 | 0.88 |
| Species-Specific OCRs and Non-OCRs | 0.43 | 0.67 | 0.66 | 0.60 | 0.73 |
| House Mouse Liver OCRs | 0.26 | 0.79 | 0.90 | 0.75 | 0.92 |
| Rhesus Macaque Liver OCRs | 0.20 | 0.84 | 0.87 | 0.77 | 0.91 |
| Norway Rat Liver OCRs | 0.45 | 0.72 | 0.90 | 0.85 | 0.79 |

| | | | | | |
|---|---|---|---|---|---|
| House Mouse Striatum OCRs | 0.45 | 0.77 | 0.63 | 0.63 | 0.77 |
| Rhesus Macaque Striatum OCRs | 0.38 | 0.79 | 0.59 | 0.76 | 0.63 |
| Norway Rat Striatum OCRs | 0.12 | 0.70 | 0.56 | 0.82 | 0.39 |
| Egyptian Fruit Bat Striatum OCRs | 0.46 | 0.83 | 0.60 | 0.71 | 0.75 |

**Table S1: Test set sensitivities, specificities, precisions, and NPVs for all evaluation criteria for house mouse-only motor cortex CNN**

| Evaluation Subset of Test Chromosome Orthologs | Fraction of Minority Class | Sensitivity | Specificity | Precision | NPV |
|---|---|---|---|---|---|
| Full Test Set | 0.47 | 0.85 | 0.81 | 0.83 | 0.83 |
| Glires-Specific OCRs and Non-OCRs | 0.061 | 0.85 | 0.89 | 0.94 | 0.71 |
| Euarchonta-Specific OCRs and Non-OCRs | 0.49 | 0.85 | 0.82 | 0.83 | 0.84 |
| Laurasiatheria-Specific OCRs and Non-OCRs | 0.36 | 0.88 | 0.84 | 0.75 | 0.92 |

| | | | | | |
|---|---|---|---|---|---|
| Species-Specific OCRs and Non-OCRs | 0.50 | 0.84 | 0.68 | 0.73 | 0.81 |
| Liver OCRs | 0.35 | 0.85 | 0.87 | 0.77 | 0.91 |
| Striatum OCRs | 0.44 | 0.85 | 0.65 | 0.76 | 0.77 |

**Table S2: Test set sensitivities, specificities, precisions, and NPVs for all evaluation criteria for multi-species motor cortex CNN**

| Evaluation Subset of Test Chromosome Orthologs | Fraction of Minority Class | Sensitivity | Specificity | Precision | NPV |
|---|---|---|---|---|---|
| Full Test Set | 0.40 | 0.75 | 0.85 | 0.77 | 0.83 |
| Glires-Specific OCRs and Non-OCRs | 0.28 | 0.69 | 0.83 | 0.91 | 0.51 |
| Euarchonta-Specific OCRs and Non-OCRs | 0.14 | 0.63 | 0.80 | 0.95 | 0.27 |
| Laurasiatheria-Specific OCRs and Non-OCRs | 0.47 | 0.74 | 0.83 | 0.80 | 0.78 |
| Species-Specific OCRs and Non-OCRs | 0.50 | 0.73 | 0.76 | 0.75 | 0.74 |
| Motor Cortex OCRs | 0.19 | 0.79 | 0.92 | 0.71 | 0.95 |

**Table S3: Test set sensitivities, specificities, precisions, and NPVs for all evaluation criteria for multi-species liver CNN**

| Evaluation Subset of Test Chromosome Orthologs | Fraction of Minority Class | Sensitivity | Specificity | Precision | NPV |
|---|---|---|---|---|---|
| Full Test Set | 0.18 | 0.64 | 0.88 | 0.54 | 0.92 |
| Full Test Set from Non-Mouse Species | 0.13 | 0.75 | 0.64 | 0.23 | 0.95 |
| Glires-Specific OCRs and Non-OCRs | 0.022 | 0.68 | 0.65 | 0.99 | 0.04 |
| Euarchonta-Specific OCRs and Non-OCRs | 0.022 | 0.69 | 0.90 | 0.14 | 0.99 |
| House Mouse Liver OCRs | 0.35 | 0.80 | 0.78 | 0.66 | 0.88 |
| House Mouse Excitatory Neuron OCRs | 0.47 | 0.70 | 0.77 | 0.73 | 0.74 |
| Human Excitatory Neuron OCRs | 0.27 | 0.82 | 0.43 | 0.79 | 0.46 |

**Table S4: Test set sensitivities, specificities, precisions, and NPVs for all evaluation criteria for house mouse-only PV+ interneuron CNN**

| Evaluation Subset of Test Chromosome Orthologs | Fraction of Minority Class | Sensitivity | Specificity | Precision | NPV |
|---|---|---|---|---|---|
| Full Test Set | 0.20 | 0.92 | 0.71 | 0.79 | 0.55 |
| Full Test Set from Non-Mouse Species | 0.20 | 0.93 | 0.73 | 0.76 | 0.54 |
| Glires-Specific OCRs and Non-OCRs | 0.49 | 0.69 | 0.74 | 0.87 | 0.45 |
| Euarchonta-Specific OCRs and Non-OCRs | 0.49 | 0.56 | 0.57 | 0.62 | 0.58 |
| House Mouse Liver OCRs | 0.32 | 0.69 | 0.56 | 0.83 | 0.61 |

**Table S5: Test set sensitivities, specificities, precisions, and NPVs for all evaluation criteria for house mouse-only retina CNN**

| Evaluation Subset of Test Chromosome Orthologs | Fraction of Minority Class | Sensitivity | Specificity | Precision | NPV |
|---|---|---|---|---|---|
| Full Test Set | 0.12 | 0.71 | 0.87 | 0.42 | 0.96 |

| | | | | | |
|---|---|---|---|---|---|
| Glires-Specific OCRs and Non-OCRs | 0.022 | 0.73 | 0.68 | 0.99 | 0.053 |
| Euarchonta-Specific OCRs and Non-OCRs | 0.022 | 0.72 | 0.87 | 0.12 | 0.99 |
| Liver OCRs | 0.35 | 0.86 | 0.78 | 0.68 | 0.91 |
| Excitatory Neuron OCRs | 0.47 | 0.74 | 0.79 | 0.76 | 0.77 |

**Table S6: Test set sensitivities, specificities, precisions, and NPVs for all evaluation criteria for multi-species PV+ interneuron CNN**

| Evaluation Subset of Test Chromosome Orthologs | Fraction of Minority Class | Sensitivity | Specificity | Precision | NPV |
|---|---|---|---|---|---|
| Full Test Set | 0.20 | 0.96 | 0.82 | 0.73 | 0.50 |
| Glires-Specific OCRs and Non-OCRs | 0.49 | 0.72 | 0.74 | 0.75 | 0.44 |
| Euarchonta-Specific OCRs and Non-OCRs | 0.49 | 0.66 | 0.66 | 0.70 | 0.50 |
| Liver OCRs | 0.26 | 0.78 | 0.90 | 0.72 | 0.26 |

**Table S7: Test set sensitivities, specificities, precisions, and NPVs for all evaluation criteria for multi-species retina CNN**

| OCR Genomic Coordinates | Permulations p-Value | phylolm Coefficient | Nearby Gene(s) of Interest [Distance to OCR ortholog] | Human Cortex Hi-C p-Value |
|---|---|---|---|---|
| chr3:6221716-62217897 (mm10) | 0.0270 | 0.624 | Arhgef26 [120] (*176*, *177*), Dhx36 [255] (*178*) | N/A, N/A |
| chr8:112950779-112951608 (mm10) | 0.0270 | 1.057 | *Cntnap4* [177] *(179, 180)*, ***Kars*** [939] (*181*, *182*) | 1, 0.9490 |
| chr18:81802310-81802951 (mm10) | 0.0270 | 0.596 | Mbp (*54*) [672], Sall3 [816] (*57*, *58*) | 1, **$2.3 \times 10^{-11}$** |
| chr16:86223026-86223725 (mm10) | 0.0270 | 0.726 | Adamts5 [322] (*183*) | **$1.8 \times 10^{-4}$** |
| chr14:78701869-78702749 (rheMac8) | 0.0270 | -0.584 | PICALM [46] (*184*), ***EED*** [128] (*185*, *186*), ***HIKESHI*** [185] | |

| | | | | |
|---|---|---|---|---|
| | | | (*187*, *188*) | |
| chr2:65033150-65033765 (rheMac8) | 0.0270 | 0.818 | CNTN6 [160] and CNTN4 [537] (*189*–*192*) | 1, 1 |
| chr10:77283085-77283984 (rheMac8) | 0.0270 | -0.473 | *SYT1* [476] (*193*) | |
| chr12:44181195-44181897 (mm10) | 0.0270 | 0.522 | *Pnpla8* [39] (*194*), Nrcam [147] (*194*) | N/A, N/A |
| chr3:105280903-105281510 (mm10) | 0.0270 | 0.482 | *Kcnd3* [171] (*195*), Mov10 [462] (*196*), <u>Lrig2</u> [769] (*197*) | 1, 1, 1 |
| chr9:15169775-15170061 (mm10) | 0.0270 | -0.438 | Vstm5 [69] (*198*), ***Med17*** [100] (*199*), *Mre11a* [373] (*200*), Piwil4 [429] (*201*), <u>Mtnr1b</u> [704] (*202*) | |

| | | | | |
|---|---|---|---|---|
| PVIL01010555:917-1512 (RouAeg_v1_BIUU) | 0.0270 | -0.592 | Hnrnpa3 [323] (*203*), *Prkra* [655] (*204*) | |
| chr2:174466184-174466517 (rheMac8) | 0.0304 | -0.575 | ***SATB1*** [509] (*48*) | |
| PVIL01006912:19061-19778 (RouAeg_v1_BIUU) | 0.0304 | -0.690 | *NEFL* (*205*, *206*) [110], ADAMDEC1 [662] (*207*), EBF2 [820] (*208*) | |
| chr12:5590611055907081 (mm10) | 0.0353 | 0.862 | *Ralgapa1* [85] (*209*), *Ppp2r3c* [603] (*210*), *Nkx2-1* [628] (*211*), Nkx2-9 [706] (*212*), Baz1a [892] (*213*, *214*) | 1, 1, 0.2746, N/A, 1 |
| chrX:69197762-69198321 (rn6) | 0.0371 | 0.578 | *Pja1* [19] (*215*), *Efnb1* [313] (*216*, *217*), *Ophn1* [560] (*218*) | N/A, N/A, N/A |

| | | | | |
|---|---|---|---|---|
| PVIL01002264:162959-163787 (RouAeg_v1_BIUU) | 0.0371 | 0.498 | *ASXL1* [26] *(219)*, KIF3B [179] (*220*), PLAGL2 [249] (*221*) | 1, 1, 1 |
| chr3:133754506-133755310 (mm10) | 0.0375 | 0.703 | ***Tbck*** [968] (*222*, *223*) | 1 |
| chr15:23008509-23009307 (rheMac8) | 0.0375 | -0.472 | ASTN2 [88] (*224*, *225*), *TRIM32* [335] (*226*, *227*) | |
| chr17:39108166-39108723 (rheMac8) | 0.0375 | 0.473 | *DIAPH3* [298] (*228*, *229*) | 1 |
| chr10:59551932-59552896 (rheMac8) | 0.0375 | 0.740 | ***USP18*** [891] (*230*) | 1 |
| chr15:105674806-105675179 (rn6) | 0.0375 | 0.729 | | |
| chrX:39249986-39250820 (mm10) | 0.0375 | 0.892 | ***Cul4b*** [674] (*231*) | N/A |
| chr12:51149370-51149713 (mm10) | 0.0375 | 0.402 | | |

| | | | | |
|---|---|---|---|---|
| chr3:140536785-140537192 (rheMac8) | 0.0375 | -0.753 | *FOXP2* [313] (*232*) | |
| chr13:101623869-101624410 (mm10) | 0.0375 | 0.485 | | |
| chr10:48660711-48661679 (rheMac8) | 0.0375 | 0.451 | MACROD2 [309] (*233*) | 1 |
| chr15:37041420-37042150 (rheMac8) | 0.0375 | 0.508 | | |
| chr1:2341557-2342331 (rn6) | 0.0375 | 1.476 | | |
| chr1:153277656-153277899 (rn6) | 0.0375 | -0.547 | *Tenm4* [433] *(234)* | |
| chr15:40082805-40083380 (mm10) | 0.0383 | 1.004 | *Dpys* [225] (*235*), *Rims2* [884] (*236*), Cthrc1 [999] (*237*) | **0.0185,** 1, 0.0640 |
| cChr2:75345159-75346046 (rheMac8) | 0.0383 | -0.654 | LRIG1 [246] (*52*, *53*) | |
| chr17:52351209-52351928 (mm10) | 0.0433 | -1.084 | ***Satb1*** [518] *(48)* | |
| chr6:28989382-28989884 (mm10) | 0.0465 | 0.801 | *Rbm28* [141] (*238*), *Smo* [746] | N/A, N/A |

| | | | |
|---|---|---|---|
| | | | (*239–241*) |
| PVIL01000147:201778-202508<br>(RouAeg_v1_BIUU) | 0.0465 | -0.570 | *TWIST1* [297]<br>(*242–244*) |

**Table S8**: **Motor cortex OCRs whose predicted open chromatin in boreoeutherian mammal orthologs is significantly associated with brain size residual after FDR correction**

Coordinates reported are those of the OCR in the species in which it was originally identified. The coefficients shown are those reported by phylolm for the association between brain size residual and OCR orthologs' predicted open chromatin. Nearby genes are those involved in brain development whose TSSs are within 1Mb of each OCR; of these, genes for which mutations cause a neurological phenotype (*47*) are in *italics* and genes for which mutations cause macro- or microcephaly specifically are in ***bold italics***. Genes important to the progression of brain cancer are underlined. Distance from the closest TSS to the ortholog of the OCR in human (gene names in all capitals) or mouse (gene names with the first letter capitalized) are reported in kb after each gene in brackets (**Materials and Methods**). The final column shows the adjusted p-value of an association in 3D space between the human ortholog of the OCR and each reported nearby gene, in order. This was done for only OCRs with predicted activity that is positively associated with brain size, since humans have the largest brain size residuals and open chromatin tends to be associated with activation (*245*). For some OCRs and genes, it was not possible to assess the association (**Materials and Methods**); those are marked with N/A. Significant adjusted permulations p-values (< 0.05) are bolded.

| OCR Genomic Coordinates | Permulations p-Value | phylolm Coefficient | Nearby Gene(s) of Interest [Distance to OCR |
|---|---|---|---|

| | | | |
|---|---|---|---|
| | | | ortholog] |
| chr1:187378056-187378556 (mm10) | 0.0117 | -1.310 | |
| chr14:12628395-12628895 (mm10) | 0.0117 | 0.490 | Fezf2 [280] (*246*), Ptprg [410] (*247*) |
| chr8:45236522-45237022 (mm10) | 0.0117 | -1.287 | Fat1 [192] (*248*), Sorbs2 [271] (*249*) |
| chr1:95762160-95762660 (mm10) | 0.0140 | -1.004 | St8sia4 [95] (*67*, *68*) |
| chr13:114793237-114793737 (mm10) | 0.0140 | -0.865 | ***Mocs2*** [25] (*66*), ***Ndufs4*** [405] (*250*, *251*) |
| chr13:36174920-36175420 (mm10) | 0.0175 | 0.490 | *Fars2* [45] (*252*), Cdyl [412] (*253*), Nrn1 [555] (*254*) |
| chr2:134681274-134681774 (mm10) | 0.0200 | -0.401 | *Plcb1* [105] (*255*), Plcb4 [977] (*256*) |
| chr16:30188019-30188519 (mm10) | 0.0219 | -1.414 | Hes1 [122] (*257*, *258*) |
| chr2:94470700-94471200 (mm10) | 0.0234 | -0.779 | ***Ext2*** [648] (*259–261*), *Alx4* [829] (*262*) |
| chr13:114757413-114757913 (mm10) | 0.0315 | -0.771 | ***Mocs2*** [61] (*66*), ***Ndufs4*** [369] (*250*, *251*) |
| chr18:25538173-25538673 (mm10) | 0.0414 | -1.005 | *Elp2* [934] (*263*) |

| | | | |
|---|---|---|---|
| chr16:30019964-30020464 (mm10) | 0.0485 | -0.901 | Hes1 [44] (*257*, *258*) |
| chr2:72557440-72557940 (mm10) | 0.0485 | -0.422 | Cdca7 [73] (*264*), Pdk1 [670] (*265*) |

**Table S9: PV+ interneuron OCRs whose predicted open chromatin in Euarchontoglires orthologs is significantly associated with brain size residual after FDR correction**

This table is in the same format as **Table S8**, except that no Hi-C p-values are included.

| Number (from Left) | Superorder | Clades(s) | Species Shown |
|---|---|---|---|
| 1 | Laurasiatheria | Eulipotyphla | *Solenodon cubanus* |
| 2 | Laurasiatheria | Ferae | *Canis latrans* |
| 3 | Laurasiatheria | Perissodactyla | *Equus africanus* |
| 4 | Laurasiatheria | Tylopoda | *Camelus dromedarius* |
| 5 | Laurasiatheria | Suina | *Sus scrofa* |
| 6 | Laurasiatheria | Ruminantia | *Bos taurus* |
| 7 | Laurasiatheria | Cetacea | *Delphinapterus leucas* |
| 8 | Laurasiatheria | Chiroptera | *Cynopterus sphinx* |
| 9 | Euarchontoglires | Lagomorpha | *Lepus americanus* |

| | | | |
|---|---|---|---|
| 10 | Euarchontoglires | Sciuromorpha + Hystricomorpha | *Hystrix sp.* |
| 11 | Euarchontoglires | Castorimorpha + Myomorpha | *Mus musculus* |
| 12 | Euarchontoglires | Strepsirrhini | *Eulemur collaris* |
| 13 | Euarchontoglires | Platyrrhini | *Callithrix jacchus* |
| 14 | Euarchontoglires | Cercopithecoidea | *Macaca mulatta* |
| 15 | Euarchontoglires | Hominoidea | *Homo sapiens sapiens* |
| 16 | Euarchontoglires | Scandentia | *Ptilocercus lowii* |

**Table S10**: **List of clades and example species shown in Figs. 3, 4, 5, and S8**

Only the Euarchontoglires clades were shown in **Figs. 4** and **5**. The Scandentia clade (16) was used in only **Fig. 5**.

| Phenotype | Ortholog threshold |
|---|---|
| Brain size | 79 Boreoeutherians |
| Solitary, group living | 42 Euarchontoglires |
| Vocal learning | 91 Boreoeutherians |

**Table S11: Thresholds for the minimum number of OCR orthologs used for each phenotype**

| OCR Genomic Coordinates | Permulations p-Value | phylolm Coefficient |
| --- | --- | --- |
| chr10:125160882-125161659 (mm10) | 0.0088 | -0.793 |
| chrX:158924956-158925400 (mm10) | 0.0088 | 0.660 |
| chr15:32784261-32784491 (rn6) | 0.0088 | -0.613 |
| chr2:72273436-72274430 (rn6) | 0.0088 | 0.790 |
| chr14:14746317-14746634 (rheMac8) | 0.0088 | 0.619 |
| chr14:10122763-10123202 (Btau_5) | 0.0088 | 0.830 |
| chr2:1375282-1375544 (Btau_5) | 0.0088 | 0.592 |
| chr:71899223-71899629 (mm1) | 0.0124 | 1.282 |
| chr18:70184720-70185120 (mm10) | 0.0124 | 0.593 |
| chr4:55604704-55605540 (mm10) | 0.0124 | 0.863 |
| chr2:7267129-7267658 (mm10) | 0.0124 | 1.552 |
| chr20:11275038-11275328 (rheMac8) | 0.0124 | 0.704 |
| chr16:3252427-3252690 (Btau_5) | 0.0124 | 0.622 |
| chr17:42269807-42270086 (Sscrofa11) | 0.0124 | 0.517 |
| chr1:257837500-257837816 (Sscrofa11) | 0.0124 | 0.878 |
| chr20:11273758-11274191 (rheMac8) | 0.0145 | 0.516 |

| | | |
|---|---|---|
| chr20:2728050-2728525 (rheMac8) | 0.0145 | 0.634 |
| chr1:13236080-13236530 (mm10) | 0.0216 | 1.218 |
| chr8:117475030-117475257 (mm10) | 0.0216 | 0.600 |
| chr19:62830511-62831002 (Btau_5) | 0.0216 | 0.528 |
| chr13:67526976-67527425 (rheMac8) | 0.0235 | 0.474 |
| chr10:95773425-95773985 (mm10) | 0.0296 | 0.730 |
| chr10:51404718-51405642 (mm10) | 0.0296 | 1.188 |
| chr1:185246181-185246600 (rheMac8) | 0.0297 | 0.473 |
| chr17:93077146-93077722 (rheMac8) | 0.0297 | 0.496 |
| chr17:93843441-93844003 (rheMac8) | 0.0298 | 0.291 |
| chr8:22552446-22552685 (Btau_5) | 0.0298 | 0.689 |
| chr15:59784092-59784782 (mm10) | 0.0298 | 3.010 |
| chr15:92199975-92200594 (rheMac8) | 0.0298 | 0.639 |
| chr13:146169647-146169932 (Sscrofa11) | 0.0309 | -0.640 |
| chr18:82975965-82976411 (mm10) | 0.0319 | 0.625 |
| chr10:17869846-17870642 (mm10) | 0.0353 | -1.045 |
| chr6:71313902-71314140 (Btau_5) | 0.0353 | -1.070 |

| | | |
|---|---|---|
| chr5:107973377-107973982 (Sscrofa11) | 0.0353 | 1.202 |
| chr7:41896644-41897066 (Sscrofa11) | 0.0353 | 0.553 |
| chr7:68829761-68830337 (mm10) | 0.0368 | 0.545 |
| chr3:182157498-182158528 (rheMac8) | 0.0368 | 0.676 |
| Chr9: (39850184-39850775 (Btau_5) | 0.0374 | 0.880 |
| chr27:22806582-22806908 (Btau_5) | 0.0380 | 0.743 |
| chr10:31826806-31827742 (mm10) | 0.0412 | 1.024 |
| chrX:33929112-33929493 (rn6) | 0.0412 | 0.624 |
| chr22:29519930-29520304 (Btau_5) | 0.0412 | 0.756 |
| chr2:200506400-200507292 (rheMac8) | 0.0431 | 0.691 |
| chr15:103102066-103102831 (mm10) | 0.0453 | 0.470 |
| chr17:85184214-85184942 (rheMac8) | 0.0453 | -0.496 |
| chr14:25533168-25533435 (mm10) | 0.0464 | 0.499 |
| chr12:18747008-18748580 (rheMac8) | 0.0464 | 0.649 |
| chr14:22931445-22931900 (rheMac8) | 0.0464 | -0.441 |
| chr13:87701407-87701687 (rheMac8) | 0.0464 | 0.481 |
| chr6:46017404-46017732 (Btau_5) | 0.0464 | 0.621 |

| | | |
|---|---|---|
| chr15:63841853-63842637 (Btau_5) | 0.0464 | 0.434 |
| chr27:3519674-3520200 (Btau_5) | 0.0464 | 0.924 |
| chr16:21486815-21487209 (rheMac8) | 0.0467 | 0.714 |
| chr6:78223211-78224022 (Sscrofa11) | 0.0469 | 0.599 |
| chr17:10702738-10703220 (mm10) | 0.0471 | 0.669 |
| chr4:40828866-40829408 (mm10) | 0.0471 | 0.523 |
| chr16:81138073-81138917 (mm10) | 0.0471 | 0.568 |
| chr6:16797433-16798215 (mm10) | 0.0471 | 0.631 |
| chr7:70625390-70625949 (mm10) | 0.0471 | 0.419 |
| chr5:24960741-24961163 (mm10) | 0.0471 | 0.493 |
| chr20:25280159-25280896 (rheMac8) | 0.0471 | 0.651 |
| chr15:87186280-87186550 (rheMac8) | 0.0471 | 0.550 |
| chr7:75139933-75140362 (rheMac8) | 0.0471 | 0.541 |
| chr9:76301902-76302190 (Btau_5) | 0.0471 | 0.602 |
| chr27:36848687-36848974 (Btau_5) | 0.0471 | 1.169 |
| chr23:48480687-48481103 (Btau_5) | 0.0471 | 0.476 |
| chr7:89298584-89298873 (Sscrofa11) | 0.0471 | -0.385 |

| | | |
|---|---|---|
| chr6:35101550-35102008 (rheMac8) | 0.0486 | 0.518 |
| chr13:43533297-43534088 (rheMac8) | 0.0486 | -0.546 |
| chr7:110183066-110183429 (Btau_5) | 0.0486 | 1.134 |
| chr21:23725264-23725522 (Btau_5) | 0.0488 | 0.465 |
| chr27:41665654-41666338 (Btau_5) | 0.0491 | 0.641 |
| chr10:81136162-81136691 (Btau_5) | 0.0491 | 0.526 |

**Table S12: Liver OCRs whose predicted open chromatin in boreoeutherian mammal orthologs is significantly associated with brain size residual after FDR correction**

Coordinates reported are those of the OCR in the species in which it was originally identified. The coefficients shown are those reported by phylolm for the correlation between brain size residual and OCR's orthologs' predicted open chromatin.
